# Reduced late endosome/lysosome function promotes SLE through chronic PI3k activity and SHP-1/SHIP-1 defects

**DOI:** 10.64898/2026.01.21.699761

**Authors:** SunAh Kang, Andrew J. Monteith, Liubov Arbeeva, Karissa Grier, Shruti Saxena-Beem, Anthony C Trujillo, Xinyun Bi, Kai Sun, Rebecca E. Sadun, Mithu Maheswaranathan, Megan EB Clowse, Saira Z Sheikh, Jennifer L Rogers, Barbara J Vilen

**Affiliations:** Department of Microbiology and Immunology, University of North Carolina; Chapel Hill, North Carolina, USA; Department of Microbiology, University of Tennessee, Knoxville, TN, USA; Division of Rheumatology, Allergy, and Immunology, Thurston Arthritis Research Center, University of North Carolina; Chapel Hill, NC, USA; Division of Rheumatology and Immunology, Duke University Medical Center; Durham, NC, USA; Lineberger Comprehensive Cancer Center, University of North Carolina; Chapel Hill, NC, USA

## Abstract

Degradation of cellular waste from phagocytosis, endocytosis and autophagy occurs through hydrolases that become activated during acidification of late endosomes and lysosomes (LELs). In a cross-sectional study we show diminished LEL acidification and the accumulation of surface-bound nucleosome on monocytes, dendritic cells, and B cells from SLE patients. Diminished acidification and exocytosis of undegraded IgG-ICs is evident in active, but not inactive disease. This is supported by our murine study where LEL acidification is diminished, promoting exocytosis and the accumulation of cell surface IgG-immune complexes. Mechanistically, LEL dysfunction is induced by chronic PI3k activation in lupus-prone MRL/*lpr* mice. We also show that on a non-autoimmune C57BL/6 background, deficiency in SHP-1 and inhibition of SHIP-1 activity is sufficient to recapitulate LEL dysfunction found in MRL/*lpr* mice. Non-acidic LELs are evident in 67% of patients, and associate with SLEDAI arthritis, rash, and nephritis. The high frequency of LEL dysfunction in SLE suggests it could serve as a biomarker identifying a specific disease endotype.

## Introduction

Systemic lupus erythematosus (SLE) is a multi-organ autoimmune disease with underlying genetic, epigenetic, and environmental components. Genome-wide association studies (GWAS) studies identified >80 confirmed risk loci (1), suggesting widespread allelic heterogeneity. Loci include genes related to lymphocyte activation, clearance of immune complexes (ICs), nucleic acid sensing, and interferon (IFN) signaling (2). Although recent therapeutic advances, including targeted biologics, have enhanced treatment options in SLE, strategies to limit the frequency and severity of active disease (3) remain challenging because factors triggering active disease remain undefined.

IgG-immune complexes (IgG-ICs) form when IgG autoantibodies bind cell-derived self-antigens, including those exposed on apoptotic blebs. IgG-ICs are associated with heightened Type I IFN responses via FcγR activation on plasmacytoid dendritic cells (pDCs), and they correlate with enhanced disease activity and lupus nephritis (4). Early studies showed that bone marrow cells from SLE patients contained undegraded, intracellular apoptotic material (5), and were termed ‘lupus erythematosus cells’. However, whether the undegraded material was due to abnormal late endosome/lysosome (LEL) function was not investigated. Later, studies of murine and human SLE identified several possible mechanisms that might increase IgG-ICs. These included heightened cell death (6), diminished phagocytosis (7), and defects in opsonins that tag apoptotic debris for removal (8).

Late endosomes/lysosomes (LELs) maintain cellular homeostasis by degrading macromolecules entering cells by endocytosis, phagocytosis, and autophagy (9). The luminal pH of endosomes/lysosomes is maintained between 4.5 to 6.5 (compared to cytoplasmic pH of ∼7.0) due to the activity of an ATP-dependent proton pump present in LEL membranes (10). Acidification of LELs facilitates release of ligands from internalized receptors and activates hydrolytic enzymes that degrade cargo (11). When the pH of LELs increases, degradation is reduced, and LELs migrate from the perinuclear region of the cell to the plasma membrane (12), where they fuse and release their contents extracellularly, a process termed exocytosis (9). Similarly, when LEL degradation is diminished, cargo in the LEL that is bound by transmembrane-spanning receptors is inserted into the plasma membrane (9).

We identified that diminished LEL acidification is evident in murine SLE (13–15), reducing the degradation of inflammatory IgG-ICs, and promoting the accumulation of nuclear self-antigen on the surface of dendritic cells (DCs), macrophages (Mφs), B cells, and T cells (13). Delayed degradation of IgG-ICs in endosomes prolongs activation of Toll-like receptors (TLRs) and disrupts the integrity of the phagolysosomal membrane, allowing IgG-ICs to leak into the cytosol (15). This provides ligands for innate cytosolic sensors including AIM2 and TRIM21, leading to pyroptosis, heightened IRF7 and increased IFNα production. Mechanistically, mTORC2 diminishes maturation and acidification of LELs in Mφs via chronic activation of Fc gamma receptor (FcγR) signaling (14). A role for FcγRs in murine SLE is also supported by bone marrow chimera studies showing that expression of activating FcγRs (FcγRI, FcγRIII, FcγRIV) on hematopoietic cells, rather than kidney mesangial cells, is required for lupus nephritis (16). Further, loss of FcγRI on myeloid cells from MRL/*lpr* mice is sufficient to attenuate B cell expansion, BAFF secretion, autoantibody production, and lupus nephritis (13). Thus, cellular homeostasis is disrupted in murine lupus when IgG-ICs and chronic FcγR signaling diminish LEL acidification.

In this study, we find that LEL dysfunction is evident in multiple splenic hematopoietic cell types in mice, multiple genetically unrelated models of SLE, and in blood cells of SLE patients. We show that diminished LEL acidification associates with active, but not inactive disease. Undegraded LEL cargo accumulates on the surface of monocytes (Mo), DCs, neutrophils, T and B cells, with the highest levels on B cells. Non-acidic LELs are evident in 67% of SLE patients, indicating a high frequency in the SLE population. Further, non-acidic LELs are found in 92% patients with active lupus nephritis, 63% with SLEDAI arthritis, and 73% with SLEDAI rash. Mechanistically, LEL dysfunction in mice is induced by PI3k activation, in part coupled to FcγRI. Deletion and/or inactivation of SHP-1 and SHIP-1 in C57BL/6 mice is sufficient to recapitulate LEL dysfunction in MRL/*lpr* mice. Our discovery of an acidification defect in LELs during active SLE is important because it could be a biomarker identifying a specific lupus endotype, and/or serve as the basis for a therapeutic aimed at maintaining disease inactivity. It is noteworthy that defects in lysosome function are emerging in multiple inflammatory diseases, including neurodegenerative diseases (Parkinson’s and Alzheimer’s (17), non-alcoholic fatty liver disease (NAFLD) (18), and Lysosomal Storage Diseases (19)). Although the molecular events underlying lysosomal defects in these diseases are unique, our studies add murine and human lupus to this growing list of inflammatory diseases with inefficient removal of waste.

## Results

### Multiple hematopoietic cell types exhibit late endosome/lysosome dysfunction

Previous studies showed that bone marrow-derived Mφ (BMMφ) from MRL/*lpr* mice exhibit diminished late endosome/lysosome (LEL) acidification (15). In all cell types, peak acidification occurred at 30 min, with de-acidification beginning at 60 min (15). To assess whether impaired acidification was evident in other cell types, we compared the LEL hydrogen ion concentration ([H+]) in splenic hematopoietic cells from MRL/*lpr* and C57BL/6 (B6) mice of different ages following stimulation with IgG-ICs. [H+] is an inversely related linear readout of pH. In 9-10 wk-old mice, the LEL [H+] in CD11b+ myeloid cells was comparable in B6 and MRL/*lpr* mice (Figure 1A); however, as disease progressed, the [H+] was decreased 12.6-fold in 15-16 wks MRL/*lpr* (Figure 1A-C; see Supplemental Table 1 for fold change and pH values). B cells (Figure 1B) and DCs (Figure 1C) from MRL/*lpr* mice also showed decreased [H+] as disease progressed (B cell: 2.3-fold; 15-18 wks; DCs: 3-fold; at >18 wks). Compared to B6 mice, the [H+] was also decreased in LELs of MRL/*lpr* neutrophils (2.1-fold) and T cells (3.2-fold) at 15-16 wks (Supplemental Figure 1A and B).

**Figure 1.**
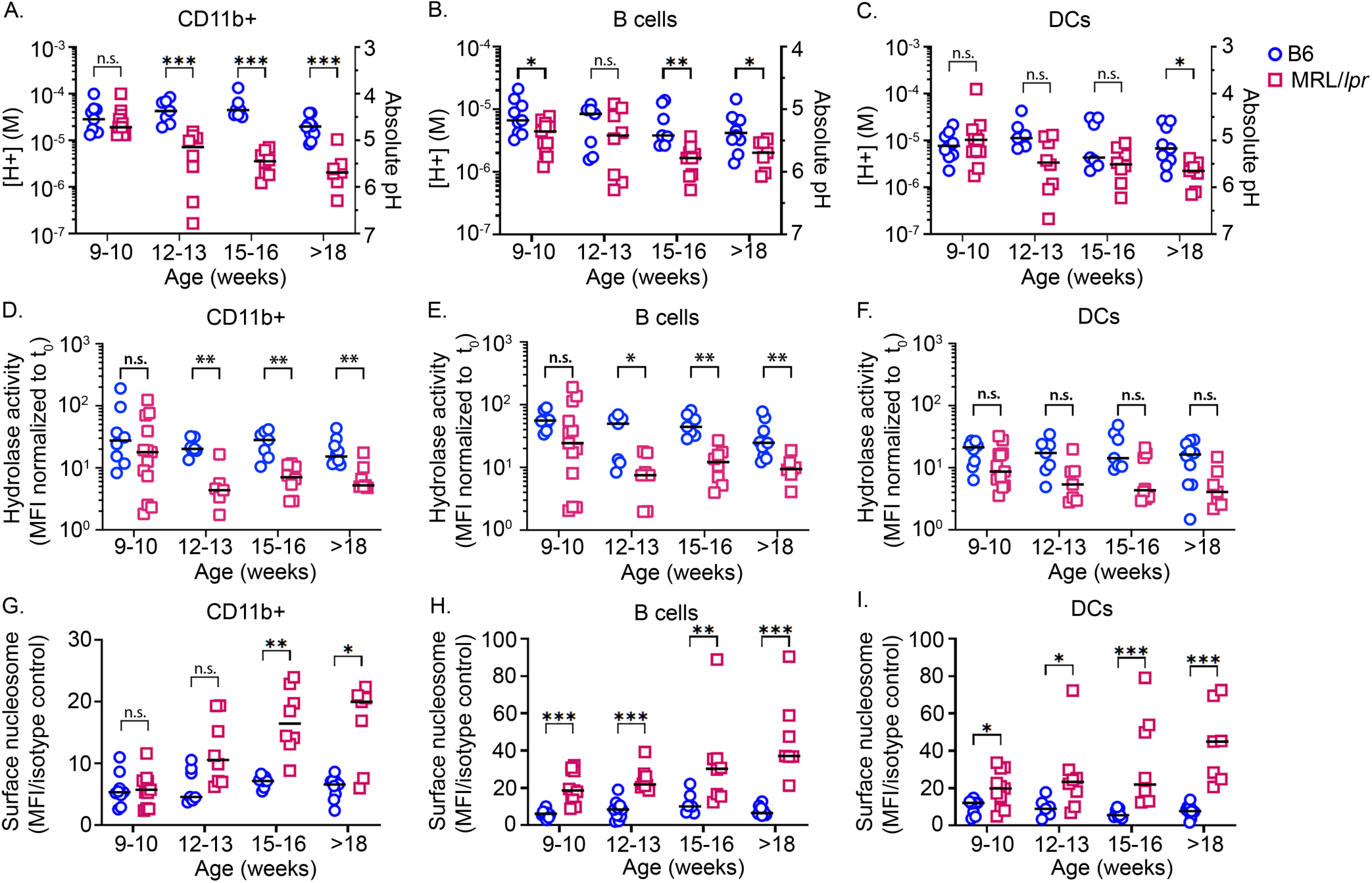
Late endosome/lysosome (LEL) dysfunction is evident in multiple hematopoietic cell types. Splenocytes from C57BL/6 (○) or MRL/*lpr* (□) mice at different ages were stimulated with IgG-ICs (30 µl IgG-ICs/0.25×10^6^ cells). LEL pH was measured by flow cytometry 30 min post-treatment, in CD11b+ myeloid cells (CD3^-^CD19^-^CD11b^+^) (**A**), B cells (CD3^-^CD19^+^) (**B**), and DCs (CD3^-^CD19^-^CD11b^+^CD11c^high^) (**C**). Absolute pH was calculated using a standard curve, then converted to [H+] (pH = -log_10_ [H+]). Flow cytometry was used to measure LEL hydrolase activity (**D-F**), and the levels of surface nucleosomes on splenocytes from mice of different ages (**G-I**). N=7-11 mice, >5 experiments per age. Statistical analysis used Mann-Whitney test (**A-I**). *p<0.05, **p<0.01, ***p<0.001. Bar = median, n.s. = not significant. See Table S1 for absolute pH and fold change calculations.

To corroborate that decreased [H+] (τpH) in LELs was biologically relevant, we quantified hydrolase activity in hematopoietic cells. The hydrolase activity in B6 CD11b+ myeloid cells (Figure 1D, 9-10 wks) was comparable to cells from MRL/*lpr* mice; however, MRL/*lpr* B cells (Figure 1E) and DCs (Figure 1F) had lower hydrolase activity compared to B6 cells (9-10 wks). Concomitant with declining [H+], hydrolase activity in CD11b+ myeloid cells and B cells from MRL/*lpr* mice was reduced 2.6 - 6.6-fold as mice aged (12- >18 wks), consistent with the idea that reduced [H+] (τpH) elicits functional consequences in LEL. Despite numerically reduced hydrolase activity in MRL/*lpr* DCs (3.2 - 4.0-fold; 12- >18 wks), the values did not achieve statistical significance (Figure 1F), suggesting that DCs maintain hydrolase activity better than myeloid cells or B cells. This might reflect that as professional antigen presenting cells (APCs), DCs predominately use cysteine proteases in late endosomes and their activation occurs at higher pH (5.0-5.5) compared to lysosomes of CD11b+ myeloid cells (20).

In MRL/*lpr* mice, diminished acidification reduces the degradation of LEL cargo (13, 15), however, cell homeostasis is maintained through exocytosis (9). To assess whether diminished LEL acidification and hydrolase activity promotes exocytosis, we quantified the levels of surface-bound nucleosome, a nuclear self-antigen in IgG-ICs. We found that splenic CD11b+ myeloid cells from B6 and MRL/*lpr* mice (9-10 wks) did not increase surface nucleosome levels, consistent with their ability to acidify and activate hydrolases; however, as MRL/*lpr* mice aged beyond 18 weeks, the surface nucleosome levels on CD11b+ myeloid cells were 3-fold higher than B6 (Figure 1G). B cells and DCs (Figure 1H and I) from 9-10 wk old MRL/*lpr* mice showed surface nucleosome levels that were 3- and 1.6-fold higher than B6, suggesting that exocytosis may occur earlier, or that these cells may have lower LEL capacity. As MRL/*lpr* mice aged beyond 18 weeks, surface nucleosome levels on B cells and DCs were increased further, to 5.8-fold compared to B6. The surface nucleosome levels on T cells and neutrophils (16-17 wk) from MRL/*lpr* mice were increased 2.1-fold (compared to B6) (Supplemental Figure 1C and D). Collectively, data from MRL/*lpr* mice show reduced LEL function in CD11b+ myeloid cells, DCs, B and T cells, and neutrophils, which worsens as mice progress to end-stage disease.

### Diminished LEL acidification is evident in genetically unrelated NZM2410 mice

To assess whether LEL dysfunction was evident in other murine lupus models, we compared B6 and MRL/*lpr* mice to MRL/MpJ, B6.*lpr*, NZM2410/J (21) and Sle123 (22). After 30 min IgG-IC stimulation, the [H+] in splenic CD11b myeloid cells from MRL/*lpr* mice was decreased 22.2-fold (p=0.0035) compared to B6 at 30min (t_30_), NZM2410/J decreased 25.3-fold (p=0.0023), B6.Sle123 16.8-fold decreased (p=0.0216), MRL/MpJ 12.6-fold decreased (p=0.0060), and B6.*lpr* showed 1.5-fold decreased (p=0.5058) (Figure 2A). LEL acidification in B6 B cells was lower than in myeloid cells; nonetheless, the [H+] in MRL/*lpr* B cells was still decreased 2.2-fold (p=0.0271), NZM2410 2.1-fold (p=0.0779), MRL/MpJ 1.9-fold (p=0.1453) and B6.*lpr* showed a 1.6-fold increase in [H+] (p=0.1428) (Figure 2B).

**Figure 2.**
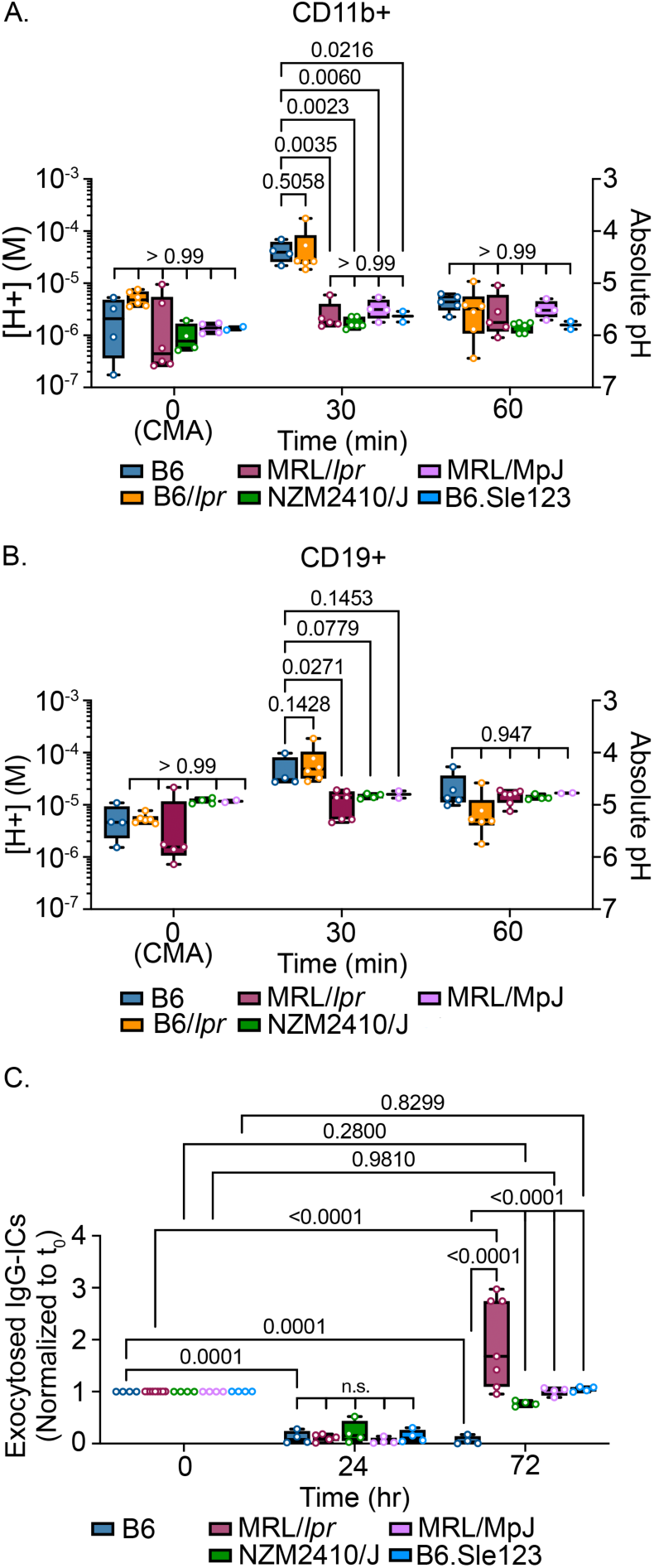
Multiple murine lupus models showed diminished [H+] and exocytosis of IgG-ICs to the plasma membrane. Splenocytes from the indicated models were stimulated with IgG-ICs (30μl IgG-ICs/0.25×10^6^ cells). At designated times, LEL pH was measured using flow cytometry in myeloid cells (CD3^-^CD19^-^CD11b^+^) (**A**) and B cells (CD3^-^CD19^+^) (**B**). vATPase activity in unstimulated samples (t_0_(CMA)), was inhibited with Concanamycin A (t_0_(CMA), 2 ng/ml). Absolute pH was calculated using a standard curve. BMMχπs were pre-loaded (t_0_) with AlexFuoro488-labeled IgG-ICs, and exocytosis was measured at designated times (**C**). Surface-bound fluorescence was assessed by subtracting internalized fluorescence (surface quenched) from total (unquenched) and normalized to individual t_0_. Statistical analysis used 2-way ANOVA with multiple comparisons (**A-C**). Adjusted *p* values with significance are shown. N = ≥2 (**A, B**) and N = ≥4 (**C**) from 2-4 separate experiments. Bar= median. Box= 25^th^-75^th^ percentiles. Whiskers= minimum and maximum values.

Exocytosis signifies undegraded cargo in LELs. To measure exocytosis, we pre-loaded BMMφ with fluorochrome-tagged IgG-ICs, establishing t_0_ levels (maximum surface fluorescence). IgG-ICs entered cells through phagocytosis, and after 24hrs incubation the surface fluorescence was reduced. Fluorescent IgG-ICs that were not degraded, returned to the cell surface via exocytosis at 72hrs (t_72_). B6 BMMφ did not undergo exocytosis, showing decreased surface fluorescence at t_72_ (35-fold decreased, p=0.0001) compared to t0. MRL/*lpr* BMMφ at t_72_ showed increased (1.7-fold, p=<0.0001) fluorochrome-tagged IgG-ICs on the cell surface compared to t_0_ levels, NZM2410 (1.3-fold, p=0.28), MRL/MpJ (1-fold, p=0.981), B6.Sle123 (1.1-fold, p=0.8299) indicative of exocytosis of undegraded LEL cargo (Figure 2C). These data show that diminished acidification is conferred by the MRL/MpJ background or the SLE123 quantitative trait loci (QTLs). The findings that diminished acidification and exocytosis are evident in genetically unrelated models of lupus raise the possibility that LEL dysfunction might be evident in human SLE.

### Active SLE patients show diminished LEL acidification and hydrolase activity

To address whether LEL dysfunction was evident in human SLE, we cross-sectionally analyzed peripheral blood cells from HCs and 81 SLE patients. Patients were grouped by disease activity (hybrid SELENA-SLEDAI; henceforth SLEDAI) as inactive (SLEDAI ≤5, n=44), moderately active (SLEDAI 6-11, n=24), or highly active (SLEDAI ≥12, n=13) (23). In our cohort (65% Black, 33% White, 9% Hispanic, 89% female, mean age of 40 yrs ± 14, mean length of disease 11 yrs ± 9.5), all ANA positive, 32% with day-of-visit renal involvement, 56% with historic renal disease, and 80% were prescribed HCQ (Supplemental Table2). As in mice, peak acidification of blood hematopoietic cells occurred at 30 min, with de-acidification beginning at 60 min. The [H+] and hydrolase activity in Mo from inactive patients were comparable to healthy controls (HC) (Figure 3A and B, Supplemental Table 3, Supplemental Figure 2). Mo from moderately active patients showed a 4.2-fold reduction in [H+] (p=0.0002), and 2-fold (p=0.04) in hydrolase activity, while highly active patients showed a 6.5-fold reduction in [H+] (p=0.0004), and 2.7-fold (p=0.007) in hydrolase activity. B cells from inactive patients showed [H+] comparable to HC (p=0.14), while hydrolase activity was decreased 2.1-fold (p=0.03). B cells from moderately active patients showed a reduction of 3.2-fold in [H+] (p=0.007) and 3.0-fold (p=0.008) in hydrolase activity, and highly active patients showed a reduction of 4.6-fold in [H+] (p=0.0009) and 3.1-fold in hydrolase (p=0.04). LELs in DCs from HCs and inactive SLE patients showed comparable [H+] (p=0.99), while hydrolase activity was decreased 2.3-fold (p=0.03) (Figure 3E and F). Compared to HC, DCs from moderately active patients showed a 5.8-fold reduction (p=0.002) with 2.9-fold decrease (p=0.01) in hydrolase activity, and highly active patients an 8.3-fold reduction in [H+] (p=0.02) with 2.4-fold decrease (p=0.01) in hydrolase activity. This shows that like murine lupus, reduced LEL acidification (↑pH) and hydrolase activity is evident in Mo, DCs and B cells from SLE patients with moderately or highly active disease, while acidification in inactive disease is comparable to HC. The exception is DCs and B cells from inactive patients, who show modestly reduced hydrolase activity. This might reflect the differences in hydrolases in late endosomes versus lysosomes, or the duration of sustained activity of hydrolases in DCs and B cells, which predominately degrade cargo in late endosome.

**Figure 3.**
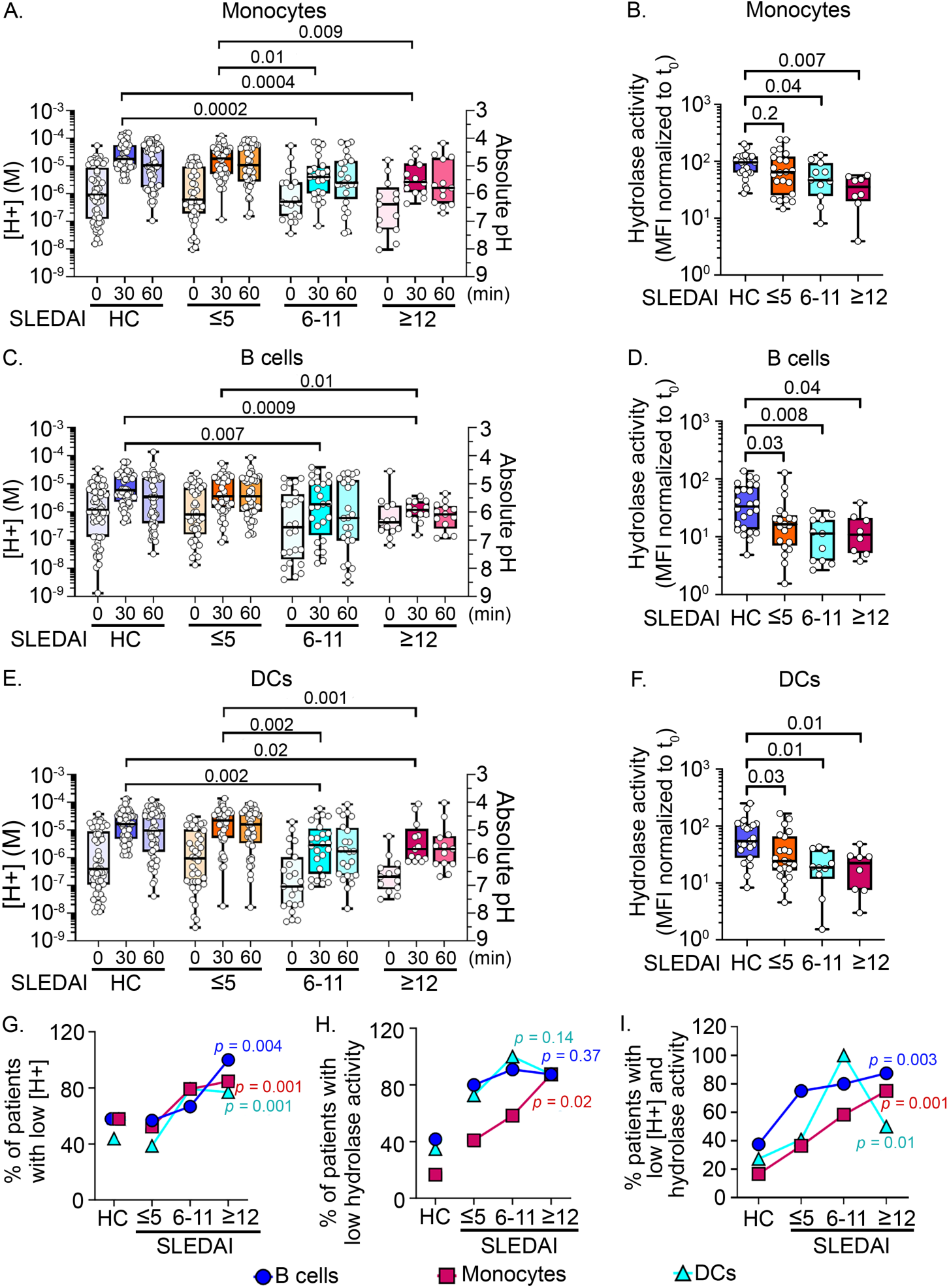
Active SLE patients show diminished LEL acidification and reduced LEL hydrolase activity. Unfractionated blood cells from HC or SLE patients were stimulated with IgG-ICs (30µl IgG-ICs/0.25×10^6^ cells). Unsimulated samples (t_0_) were treated with Concanamycin A (20ng/ml) to inhibit vATPase activity. At designated times, LEL pH was measured in each cell type (**A, C**, **E**). Absolute pH was calculated using a standard curve. The LEL hydrolase activity was measured by flow cytometry using an acidotropic substrate that fluoresces upon degradation (**B**, **D**, **F**). The hydrolase substrate MFI was normalized to t0. Trends (**G-I**) were assessed using the Cochran-Armitage test to compare the proportion of patients in each disease group with low [H+] (**G**), low hydrolase activity (**H**), or low [H+] and hydrolase activity (**I**) for B cells (●), monocytes (▪), DCs (▴). In (**A**, **C**, **E**), HC N = 57, SLE N = 81. In (**B**, **D**, **F**), HC N = 24, SLE N = 41, >8 experiments. Statistical analysis used Kruskal-Wallis (**A-F**). Adjusted *p* values with significance are shown. Bar = median. Box= 25^th^-75^th^ percentiles. Whiskers= minimum and maximum values.

To assess associations between reduced LEL [H+] and disease activity, we calculated the proportion of patients with low [H+] in each SLEDAI group (SLEDAI <5, 6-11, or ≥12), then used the Cochran-Armitage test to identify trends between the proportions. The proportion of patients with low [H+] increased as the SLEDAI groups increased in disease activity. Patients with “non-acidic” LELs have [H+] lower than a cut-off that was set at 1.8-fold above the mean [H+] of HC for each cell type. (Mo p=0.001, B cell p=0.004, DC p=0.001) (Figure 3G). These data suggest an association between SLE disease activity and diminished LEL acidification. The proportion of patients whose Mo show low hydrolase activity also increased across SLEDAI groups (p=0.02; Figure 3H); except in B cells (p=0.37) or DCs (p=0.14), likely because during inactive disease, B cells and DCs have modestly reduced hydrolase activity (Figure 3D and F). Patients with “low hydrolase” have levels below a cut-off that was established at 1.7-fold above the mean hydrolase of HC for each cell type. The proportion of patients with low [H+] and low hydrolase increased as the SLEDAI groups increased in disease activity (Mo p=0.001, B cell p=0.003, DC p=0.01) (Figure 3I). Finally, we estimated the frequency of SLE patients (regardless of disease activity) with diminished LEL acidification. In our cohort of 81 patients, 67% had non-acidic LELs in B cells, 65% in Mo, and 57% in DCs (Supplemental Table 5). These data reveal that LEL dysfunction affects a significant portion of SLE patients and suggests an association between SLEDAI groups and the efficiency of LEL function, especially in Mo.

### LEL dysfunction is not associated with hydroxychloroquine treatment

The mechanism of action of HCQ was initially described as alkalization of LELs, which reduced antigen presentation (24). However, more recent studies show that HCQ-mediated LEL alkalinization is transient, with normal pH restored within 4-hours (25). We reasoned that if HCQ was responsible for disrupting LEL acidification, then regardless of disease activity, non-acidic LELs would be more prevalent among patients prescribed HCQ, compared to those not prescribed HCQ. The data show that 82% of patients with acidic LELs, and 79% of patients with non-acidic LELs (in Mo), were prescribed HCQ. Similar results were found with B cells (74% acidic, 83% non-acidic) and DCs (83% acidic and 78% non-acidic; Table 1) Further, since non-acidic LELs associated with increased disease activity (Figure 3G), we reasoned that if HCQ caused non-acidic LELs, then the proportion of patients prescribed HCQ should increase as disease activity increased. Instead, the proportion prescribed HCQ was similar across SLEDAI groups, (82% inactive, 79% moderately active, 69%-77% highly active; Table 1). This suggests that diminished LEL acidification is not a consequence of HCQ, a finding consistent with LEL dysfunction in untreated MRL/*lpr* mice (13–15). Alternative explanations for the efficacy of HCQ in treating SLE include its ability to intercalate into nucleic acids (26) and inhibit nucleic acid binding to innate cytosolic sensors (27) and endosomal TLRs (28). HCQ also reduces reactive oxygen species (ROS) by blocking NOX2 assembly (29), Ca^2+^ release from the ER (30), and CD40L expression (31), events that decrease cellular activation and the secretion of inflammatory cytokines.

**Table 1.** Proportion of patients prescribed hydroxychloroquine (HCQ) among patients with acidic versus non-acidic late endosomes/lysosomes (LELs), or with varying disease activity for each cell type.

|  | Overall | Patients with<br>acidic LEL | Patients with<br>non-acidic<br>LEL<sup>A</sup> | SLEDAI ≤5 | SLEDAI 6-11 | SLEDAI ≥12 |
| --- | --- | --- | --- | --- | --- | --- |
| N | 81 |  |  |  |  |  |
| patients prescribed HCQ N(%) | 65 (80) |  |  |  |  |  |
| Monocytes |  |  |  |  |  |  |
| N |  | 28 | 53 | 44 | 24 | 13 |
| patients prescribed HCQ N(%) |  | 23 (82) | 42 (79) | 36 (82) | 19 (79) | 9 (69) |
| B cells |  |  |  |  |  |  |
| N |  | 27 | 54 | 44 | 24 | 13 |
| patients prescribed HCQ N(%) |  | 20 (74) | 45 (83) | 36 (82) | 19 (79) | 10 (77) |
| DCs |  |  |  |  |  |  |
| N |  | 35 | 46 | 44 | 24 | 13 |
| patients prescribed HCQ N(%) |  | 29 (83) | 36 (78) | 36 (82) | 19 (79) | 10 (77) |

<sup>A</sup>Patients with “non-acidic” LELs have [H+] lower than the cut-off. For each cell type, the [H+] cut-off was established at 1.8-fold above the mean [H+] of the HC (monocytes pH 4.7, B cells pH 5.1, DCs pH 4.8).

### Nuclear self-antigens accumulate on blood Mo, DCs, and B cells during highly active disease

To assess whether SLE patients accumulate nuclear antigens on the plasma membrane, we quantified surface nucleosome on blood cells from SLE patients and HC. In patients with inactive and moderately active disease, the levels of surface nucleosome on Mo (Figure 4A), B cells (Figure 4B), and DCs (Figure 4C), were comparable to HC. However, in highly active disease, surface nucleosome levels were increased on Mo (1.8-fold), B cells (7.1-fold), and DCs (1.7-fold). Accumulation of surface nuclear antigen on hematopoietic cells was not unique to nucleosome as highly active SLE patients showed 2.9-fold increased surface dsDNA on Mo (p=0.03) and 5.3-fold increase on B cells (p=<0.0001) (Supplemental Figure 3A and B; Supplemental Table 6). It is also possible that accumulation of nucleosome reflected increased FcγR expression, however, the levels of FcγRI, FcγRIIA, FcγRIII and FcγRIIb on Mo, DCs, and B cells from SLE patients were not different from HC (p=0.23-0.97) (Supplemental Figure 4). It is noteworthy that some of the anti-FcγRs have epitope specificity for the Fc-binding cleft (blocking antibody) and may not have detected FcγRs that were pre-bound to IgG-ICs. The broad histogram peaks reveal cell-to-cell variability in surface nucleosome, most notably on B cells in highly active disease (Figure 4B).

**Figure 4.**
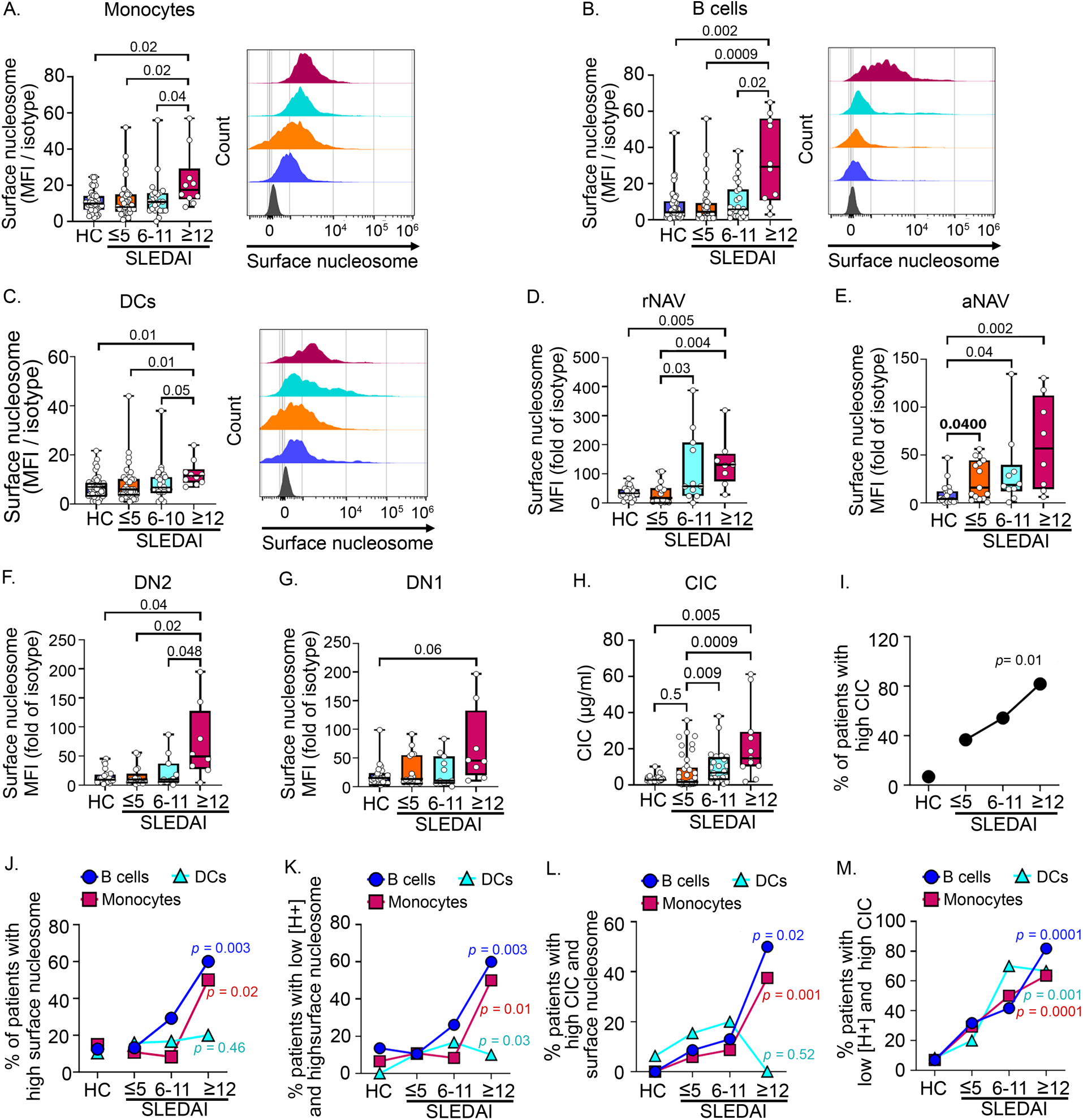
Highly active SLE patients (SLEDAI ≥12) show elevated levels of surface nucleosome and circulating immune complexes (CIC). Unfractionated blood Mo (**A**), B cells (**B**), DCs (**C**), rNAV (**D**), aNAV (**E**), DN2 (**F**), DN1 (**G**) were analyzed for surface nucleosome by flow cytometry. Representative histograms show the cell distribution with varying disease activities (gray: isotype control). Plasma CIC levels were measured by IC-mediated internalization of FcγRIIA on neutrophils using flow cytometry and a standard curve **(H).** N = 11-41 per disease group, 3 experiments. Trends (**I-M**) were assessed on B cells (●), Mo (▪), DCs (▴) using the Cochran-Armitage test to compare the proportion of patients in each disease group with high CIC **(I)**, surface nucleosome (**J**), low [H+] and high surface nucleosome (**K**), high CIC and surface nucleosome (**L**), and low [H+] and high CIC (**M**). In (**A-C**) HC N = 48, SLE patient = 72, in (**D-G**) HC N = 19, SLE patient = 8-15. Statistical analysis used Kruskal-Wallis (**A-H**). Adjusted *p* values with significance are shown. Bar = median. Box= 25^th^-75^th^ percentiles. Whiskers= minimum and maximum values.

To identify the B cell subset(s) with elevated levels of surface nucleosome, we quantified surface nucleosome on IgD^pos^CD27^neg^ resting naïve (rNAV), activated naive B cells (aNAV) and the IgD ^neg^CD27 ^neg^ double negative 1 (DN1) and 2 (DN2) B cells (Supplemental Figure 5) (32). aNAV B cell are the precursors of DN2 cells that differentiate into antibody secreting cells through extrafollicular response. The frequency of rNAV, aNAV, DN1 and DN2 B cells in SLE patient and HC blood showed a significant expansion of aNAV and DN2 cells as previously described (32). Comparing surface nucleosome levels to HC, we found levels on rNAV were increased 2.0- and 4.4-fold in moderately and highly active disease (Figure 4D), while surface nucleosome on aNAV was increased 4.0-, 4.8-, and 14.6-fold across the SLEDAI groups (Figure 4E). In highly active disease, surface nucleosome on DN2 cells were increased 4.1-fold compared to HC (Figure 4F). Although surface nucleosome was elevated on DN1 cells in highly active disease, the levels were not statistically different than HC (p=0.06, Figure 4G). Thus, during highly active disease, surface nucleosome levels were increased on all cell types; however, the aNAV B cell subset shows the highest levels.

In addition to elevating surface nucleosome, exocytosis also elevates CIC levels. We found that the CIC levels in inactive patients did not change compared to HC; however, in patients with moderately and highly active disease were increased 2.2- and 4.7-fold compared to HC (p=0.009, 0.0009) (Figure 4H, Supplemental Table 8). Patients with “high CIC” show plasma levels higher than a cut-off established at 1.5-fold above the mean CIC level of HC. The proportion of patients with elevated CIC also increased over the SLEDAI groups, revealing an association between CIC levels and disease activity (p= 0.01, Figure 4I). To identify whether increasing disease activity was related to LEL dysfunction, we used Cochran-Armitage analysis. For this analysis, patients with “high surface nucleosome” were defined as having cell levels above a cut-off established at 1.8-fold above the mean surface nucleosome level of HC. We found a significantly higher proportion of patients in highly active disease showed high surface nucleosome on Mo and B cells (Figure 4J) (Mo p=0.02, B cells p=0.003); however, this trend was not seen with DCs (p=0.46). In B cells, the trend was corroborated by a high Spearman’s correlation coefficient in patients with highly active disease (r=0.81, p=0.004, Supplemental Table7). We also found that a higher proportion of patients with active disease showed both low [H+] and elevated surface nucleosome on B cells (p=0.003), Mo (p=0.01), and DCs (p=0.03; Figure 4K), or high CIC and increased surface nucleosome on B cells (p=0.02) and Mo (p=0.001), but not DCs (p=0.52) (Figure 4L), or elevated CIC and low [H+] on B cells (p=0.0001), Mo (p=0.001), and DCs (p=0.0001) (Figure 4M). The weaker trend in DCs could reflect that exocytosis occurs after DC migrate to lymph nodes and complete their maturation (33), since CIC induce CCR7-dependent migration of DCs to lymph nodes in both human and murine lupus (34). In summary, SLEDAI groups with higher disease activity are associated with decreased hydrolase activity and non-acidic LELs in Mo and B cells, and increased surface nucleosome and CIC on Mo, B cells and DCs. These findings support the idea that LEL dysfunction associates with SLEDAI groups of higher disease activity.

### LEL dysfunction is not evident in active rheumatoid arthritis patients

To assess whether reduced LEL [H+] and the accumulation of surface nucleosome were evident in other rheumatic diseases, we analyzed blood hematopoietic cells from active, seropositive rheumatoid arthritis (RA) patients (n=23, Supplemental Table 9). The [H+] and levels of surface nucleosome on blood hematopoietic cells from active RA patients were not different compared to HC (Supplemental Figure 6). This indicates that the hallmarks of LEL dysfunction are not evident in blood cells from active RA.

### Patients with LEL dysfunction are more likely to have renal disease, rash, and arthritis

To identify relationships between clinical symptoms and LEL dysfunction, we separated patients with non-acidic LELs, then calculated the proportion of this group receiving disease-modifying anti-rheumatic drugs (DMARDs; Mycophenolic acid, Mycophenolate mofetil, Azathioprine, Methotrexate, Tacrolimus), or having clinical manifestations involving renal, skin, or joint. Of the patients with non-acidic LELs, 44% showed current renal disease, 35% SLEDAI rash, 22% SLEDAI arthritis, and 72% were receiving

DMARDs (Supplemental Table 5). In addition, we identified patients with each clinical manifestations (renal, skin, joint, or receiving DMARDs), then calculated the proportion of this subgroup with non-acidic LELs. Of those with current renal disease, rash, arthritis or receiving DMARDS, 92%, 73%, 63%, and 75% had non-acidic LELs. (Supplemental Table 4). Thus, when patients show clinical manifestations, they are more likely to already exhibit dysfunctional LELs, while patients with non-acidic LELs may not have developed clinical symptoms. This raises the possibility that diminished LEL acidification is coincident with or could precede clinical manifestations.

To identify whether LEL dysfunction associates with renal disease, we grouped SLE patients (regardless of disease activity) into those with active nephritis, remission nephritis, or those who never had renal disease (never nephritis), then compared the proportion with nonacidic LELs or increased surface nucleosome. A higher proportion of SLE patients with active renal disease had low [H+] (80-92%; mean SLEDAI of 11) compared to the proportion of patients who never had nephritis (49-57%), or remission nephritis (45-55%). This suggests that the function of LELs is restored in remission nephritis. Similarly, more patients with active nephritis showed elevated nucleosome on the surface of B cells (48%; Supplemental Table 10). These findings suggest that LEL dysfunction associates with active nephritis, is characterized by low [H+] in all cell types, and the accumulation of surface nucleosome on Mo, DCs and B cells from highly active patients. Collectively, our findings show that LEL dysfunction associates with active clinical symptoms (SLEDAI rash, arthritis, and nephritis) and is coincidental with, or could precede, these manifestations.

### FcγRI is coupled to LEL dysfunction in MRL/*lpr* mice

Consistent with a role for FcγRI in murine SLE, we previously showed that MRL/*lpr* mice lacking FcγRI (FcγRI^-/-^/MRL/*lpr*) did not develop lupus and showed diminished signaling of pSyk^Y525^, pAkt^S473^, pAkt^T308^, pS6 (13, 14). To assess whether FcγRI^-/-^/MRL/*lpr* mice restore LEL function, we used BMMφ and measured acidification ([H+]), and ROS. After 30 min stimulation with IgG-ICs, the [H+] in MRL/*lpr* BMMφ was reduced 8.3-fold (p=0.003), while in FcγRI^-/-^/MRL/*lpr* BMMφ [H+] was only reduced 1.8-fold (p=0.0845) (Figure 5A). The levels of ROS in B6 BMMφ were increased 3.8-fold (p<0.0001) at 15min, returning to 1.4-fold (p=0.288) at 2hrs (Figure 5B). In MRL/*lpr* BMMφ, ROS was increased 5.8-fold (p<0.0001) (15min) and sustained at 7.3-fold (p<0.0001) at 2hrs (compared to MRL/*lpr* t0). In FcγRI^-/-^/MRL/*lpr* BMMφ ROS was increased 2.7-fold (p=0.35) (15 min) and sustained at 2.7-fold (p=0.707) at 2hrs (compared to FcγRI^-/-^/MRL/*lpr* t_0_). Thus, loss of FcγRI in MRL/*lpr* mice reduces the peak ROS levels, but those levels are sustained over 2hs. We previously identified that FcγRI plays an important role in murine SLE (13), and now show that FcγRI is required for diminished acidification and heightened ROS in MRL/*lpr* mice, showing a contribution to LEL dysfunction.

**Figure 5.**
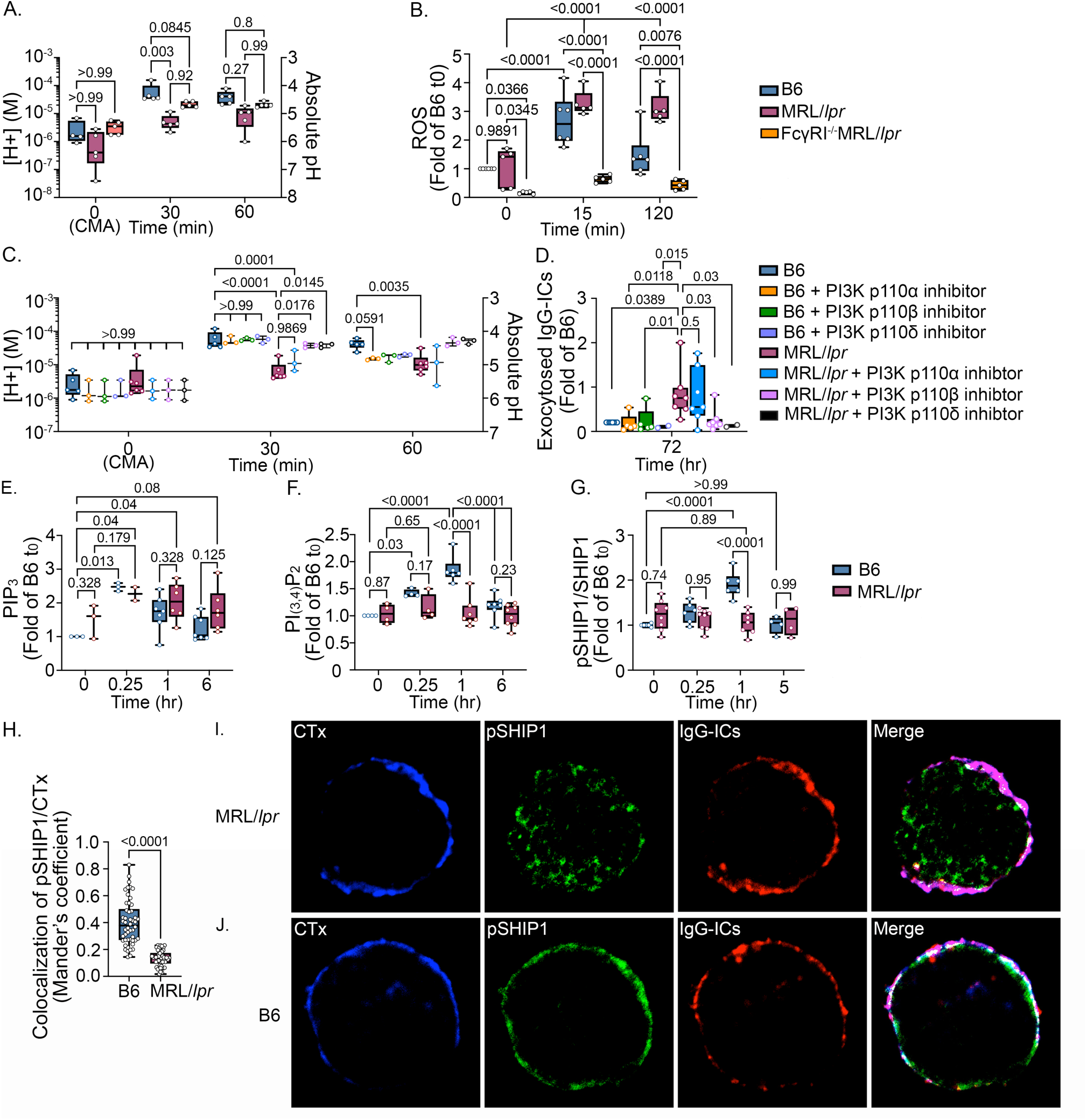
LEL defects are induced by chronic PI3K activation and SHIP-1 defects, and evident in FcγRI^-/-^MRL/*lpr* mice. BMMχπs were stimulated with IgG-ICs (25μl IgG-ICs/0.25×10^6^ cells) (**A-J**). At designated times, LEL pH (**A, C**), ROS (**B**), exocytosis (**D**), PIP_3_ (**E**), PI_(3,4)_P_2_ (**F**), pSHIP^Y1022^ (**G**) were measured by flow cytometry. vATPase activity in unstimulated samples (t_0_(CMA)) was inhibited with Concanamycin A (CMA, 2 ng/ml) (**A, C**). ROS levels (**B**) were measured using CellROX including t0 samples untreated with IgG-ICs. BMMχπs were pre-loaded (t0) with AlexFuoro488-labeled IgG-ICs, and exocytosis was measured at designated times (**C**). Surface-bound fluorescence was assessed by subtracting internalized fluorescence (surface quenched) from total (unquenched) and normalized to individual t0. The effect of PI3K was measured using PI3k-p110 inhibitors (p110α, β, and 8) (100nM, 2hrs prior to IgG-IC treatment) (**C, D**). The colocalization of IgG-ICs (red), pSHIP^Y1022^ (green) with cholera toxin-positive lipid rafts (CTx, blue) (**H**) in BMMχπs was analyzed by confocal microscopy (**I, J**). Images were processed using Image J. Representative images are shown. White in merged images depicts colocalized IgG-ICs, pSHIP^Y1022^, and CTx. Statistical analysis used 2-way ANOVA with multiple comparisons (**A-G**) and Mann-Whitney test (**H**). Adjusted *p* values with significance are shown. N= 2-8, >2 experiments (**A-H**), N = 3, 3 experiments, total of 50 cells/mouse line (**H-J**). Bar = median. Box= 25^th^-75^th^ percentiles. Whiskers= minimum and maximum values.

### Chronic PI3k activity impairs LEL function

Past studies of LEL dysfunction identified a pathway where the binding of cofilin to phagosomal actin is impaired due to heightened cofilin phosphorylation. This diminishes Rab39a cleavage (14), a necessary step in lysosomal acidification (35). Since FcγRI is coupled to the cofilin/actin pathway through PI3k/Akt signaling, we tested whether inhibiting PI3k-p110 activity restored LEL function. BMMφs from B6 and MRL/*lpr* mice were treated with isoform inhibitors of the PI3k-p110 subunit; -p110α (PIK-75), -p110β (TGX-221), or -p110δ (IC87114). In MRL/*lpr* BMMφs, the [H+] was decreased 9.2-fold (p<0.0001) compared to B6 t_30_ (Figure 5C). In MRL/*lpr* BMMφs, the -p110α inhibitor did not restore an acidic [H+], instead maintain 3.9-fold decreased [H+] (p=0.0001) compared to B6 t_30_. In contrast, treatment with inhibitors of -p110β or -p110δ increased the [H+] to levels comparable to B6 t_30_ levels (1.1-fold p=0.2503; 1.1-fold p=0.2812). Inhibiting PI3k-p110β or -p110δ, but not PI3k-p110α, in MRL/*lpr* BMMφs also prevented exocytosis (Figure 5D). The levels of fluorochrome-tagged IgG-ICs on MRL/*lpr* BMMφs, were 3.8-fold higher than untreated B6 BMMφs (p=0.0389). However, after treatment with -p110β or -p1108 inhibitors [H+] were comparable or below the levels in B6 BMMφs. In contrast, the levels of exocytosis in MRL/*lpr* BMMφs treated with the -p110α inhibitor were not different from untreated MRL/*lpr*, (p=0.5). These data demonstrate that chronic PI3k activity of -p110β and -p110δ in MRL/*lpr* BMMφs contributes to diminished LEL acidification and reduced degradation of IgG-ICs. It also corroborates previous data showing reduced lupus nephritis, B cell expansion, BAFF and autoantibody production in FcγRI^-/-^/MRL/*lpr* mice (13).

To understand how PI3k activity contributes to LEL dysfunction, we analyzed the products of PI3k activation in IgG-IC stimulated B6 and MRL/*lpr* BMMφs. PI3k activation converts PI(4,5)P2 to PI(3,4,5)P3 (PIP3) (Figure 5E). In B6 BMMφs, basal levels of PIP_3_ increased 2.5-fold (p=0.013) after 15 min stimulation, then returned to basal levels. In MRL/*lpr* BMMφs basal levels of PIP_3_ were increased 1.6-fold (compared to B6 t_0_). Following 15min stimulation, PIP3 levels further increased 1.4-fold (p=0.179) to levels that were 2.3-fold (p=0.04) higher than B6 t0. At 6hrs, PIP3 levels remained relatively high at 1.7-fold (p=0.08) compared to B6 t0. The elevated basal levels, and low IgG-IC-induced levels, of PIP3 suggests sustained PI3k activity in MRL/*lpr* mice. Alternatively, diminished phosphatase activity, could heighten PIP3. To gain insight into phosphatases regulating PIP3 levels, we examined other phosphoinositide products. Activation of SHIP-1 dephosphorylates PIP3 to produce PI(3,4)P_2_ (36). In B6 BMMφs, PI(3,4)P_2_ levels were transiently increased over 1 hr (1.8-fold, p=<0.0001), but unchanged in MRL/*lpr* (1.0-fold, p=0.79) (Figure 5F). Reduced formation of PI(3,4)P_2_ following IgG-IC stimulation of MRL/*lpr* BMMφs suggests impaired SHIP-1 activity.

To assess whether phosphorylation of SHIP-1 reflected the phosphoinositide products, we compared levels of pSHIP-1^Y1022^ in B6 and MRL/*lpr* BMMφs (Figure 5G). B6 BMMφs increased pSHIP-1^Y1022^ 1.9-fold, (p=<0.0001, compared to B6 t_0_) after 1hr stimulation which returned to baseline by 5 hrs. Unstimulated MRL/*lpr* BMMφs had slightly higher basal pSHIP-1^Y1022^ (1.25-fold, p=0.74 compared to B6) that remained unchanged over 5hrs (1.2-fold; p>0.99, compared to MRL/*lpr* t_0_). The results support the idea that impaired SHIP-1 phosphorylation could account for the sustained PIP_3_ levels in MRL/*lpr* BMMφs. It also raised the possibility that impaired phosphorylation of SHIP-1 in MRL/*lpr* BMMφs could reflect decreased recruitment of SHIP-1 to the plasma membrane to localize with FcγRIIb (36). To assess this, we used confocal microscopy to quantify the levels of pSHIP-1^Y1022^ colocalized with cholera toxin (CTx) stained membrane lipid rafts (Figure 5H-J). Following 1 hr stimulation with IgG-ICs, the levels of pSHIP-1^Y1022^ colocalized with CTx-positive lipid rafts on the plasma membrane of MRL/*lpr* BMMφs (Figure 5I) were 2.7-fold (p<0.0001) decreased compared to B6 (Figure 5J). Taken together, these results demonstrate that MRL/*lpr* BMMφs show decreased SHIP-1 phosphorylation and fail to localize pSHIP-1 to the lipid rafts and the site of FcγRI.

### Diminished SHIP-activity in non-autoimmune mice partially impairs LEL function

Src homology 2-containing inositol phosphatase-1 (SHIP-1, *Inpp5d*) is activated by phosphorylation at Y^1022^ (37) and recruited to FcγRI through ITIM-containing FcγRIIb. We hypothesized that if SHIP-1 was unable to efficiently dephosphorylate PIP3 in MRL/*lpr* BMMφs and consequently induce LEL dysfunction, then B6 BMMφs deficient in SHIP-1 (B6.SHIP-1^-/-^) should recapitulate LEL dysfunction in MRL/*lpr* mice. Following stimulation with IgG-ICs, B6.SHIP-1^-/-^ BMMφs showed a 3.7-fold decrease (p=0.01) in [H+] at 30 min, while MRL/*lpr* showed a 10.3-fold decrease (p<0.0001) (compared to B6 t0), suggesting that SHIP-1 deficiency does not fully recapitulate diminished acidification in MRL/*lpr* mice (Figure 6A). Similarly, at 60 min post-stimulation, the B6.SHIP-1^-/-^ BMMφs did not show increased [H+], confirming that delayed acidification was not occurring (data not shown). When stimulated with IgG-ICs for 15min, ROS levels in B6.SHIP-1^-/-^ BMMφs increased 3.3-fold (p=0.03) compared to B6.SHIP-1^-/-^ t_0_, MRL/*lpr* 2.1-fold (p=0.03), and B6 2.0-fold (p=0.03) (Figure 6B). At 2 hrs, ROS declined in B6.SHIP-1^-/-^ (1.6-fold, p=0.72) and B6 (1.3-fold, p=0.28) compared to their individual t_0_, while in MRL/*lpr*, ROS levels remained high (2.0-fold, p=0.04). This indicates that SHIP1 deficiency does not recapitulate the elevated levels of sustained ROS seen in MRL/*lpr* mice. We then assessed whether SHIP1 deficiency increased exocytosis of undegraded IgG-ICs (Figure 6C). MRL/*lpr* BMMφs showed slightly higher levels (1.2-fold) of fluorochrome-labeled IgG-ICs at 72hr compared preloaded levels at t_0_ (p=0.76), indicating exocytosis of IgG-ICs to plasma membrane. In contrast, B6 and B6.SHIP-1^-/-^ BMMφs showed surface fluorescence comparable or below t_0_, consistent with IgG-IC degradation. Like B6.SHIP-1^-/-^, B6 BMMφs treated with a SHIP-1 inhibitor (3AC) showed modestly decreased [H+], significantly elevated but not sustained ROS, with no exocytosis (Figure 6A-C). Together, the data show the SHIP-1 deficiency is not sufficient to recapitulate the LEL dysfunction of MRL/*lpr* mice.

**Figure 6.**
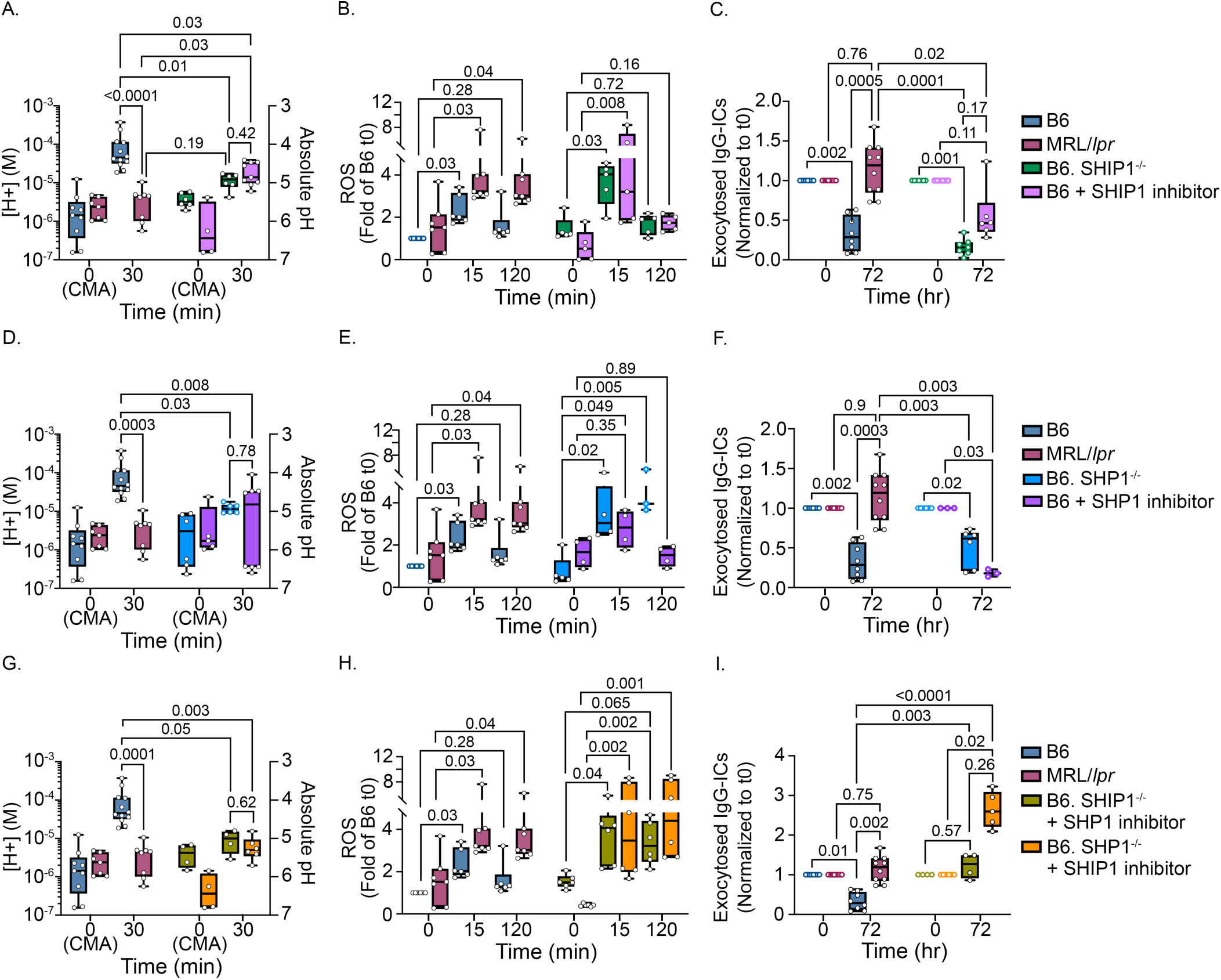
Deficiency in SHP-1 and inhibition of SHIP-1 in B6 mice phenocopies the LEL dysfunction seen in MRL/*lpr*. To assess the effects of SHIP1 and/or SHP1 on LEL defects, BMMχπs from B6, B6.SHIP1^-/-^, and B6.SHP1^-/-^ were treated ± inhibitors for SHP1 (10μM NSC-87877, 3hrs prior to IgG-IC treatment) or SHIP1 (50nM 3AC, 48hrs prior to IgG-IC treatment). The effects of single deficiency of SHIP1 (**A-C**) or SHP1 (**D-F**), or double deficiency (**G-H**) were analyzed. BMMχπ were stimulated with IgG-ICs (25μl IgG-ICs/0.25×10^6^ cells) ± inhibitors. At designated times, LEL pH (**A, D, G**), ROS (**B, E, H**), and exocytosis (**C, F, I**) were measured by flow cytometry. Absolute pH was calculated using a standard curve (**A, D, G**). vATPase activity in unstimulated samples (t_0_(CMA)) was inhibited with Concanamycin A (CMA, 2 ng/ml). ROS levels (**B, E, H**) were measured using CellROX including t0 samples untreated with IgG-ICs, and fold of B6 t_0_ was graphed. BMMχπs were pre-loaded (t0) with AlexFuoro488-labeled IgG-ICs, and exocytosis was measured at designated times (**C, F, I**). Surface-bound fluorescence was assessed by subtracting internalized fluorescence (surface quenched) from total (unquenched) and normalized to individual t0. Statistical analysis used Kruskal-Wallis test with multiple comparisons. Adjusted *p* values with significance are shown. N= 4-12, 3-4 experiments. Bar = median. Box= 25^th^-75^th^ percentiles. Whiskers= minimum and maximum values.

SHP-1 *(Ptpn6)*, a protein tyrosine phosphatase that dephosphorylates the FcγR gamma-chain (FcRγ;FCER1G) ITAM motif (YxxL/I) (38), decreases Syk recruitment (39) and dampens FcγRI signal transduction (40). If heightened PI3k activity in LEL dysfunction occurs through sustained FcγRI signaling, then B6.SHP-1-deficient mice (B6.SHP-1^fl/fl^ x B6.Rosa26-CreERT2) should recapitulate the LEL defect seen in MRL/*lpr* BMMφs. Tamoxifen treatment of SHP-1-deficient mice efficiently excised SHP-1 (Supplemental Figure 7). Following 30 min stimulation with IgG-ICs, B6.SHP-1^-/-^ BMMφs showed 3.9-fold (p=0.03) lower [H+] compared to 10.3-fold (p=0.0003) lower in MRL/*lpr* (compared to B6 t_30_, Figure 6D). After 15 min of IgG-IC stimulation, acute ROS levels in B6.SHP-1^-/-^ BMMφs were increased 6.5-fold (p=0.02) compared with a 2.1-fold (p=0.03) increase in MRL/*lpr* (Figure 6E). At 2 hrs, sustained ROS in B6.SHP-1^-/-^ BMMφs were 8.5-fold over t_0_ (p=0.005) while levels in MRL/*lpr* were sustained at 2.0-fold (p=0.04), indicating that SHP1 deficiency heightens and sustains ROS. B6.SHP-1^-/-^ BMMφs did not show exocytosis of undegraded IgG-ICs (Figure 6F). Like the B6.SHP-1^-/-^ BMMφs, B6 BMMφs treated with a SHP-1 inhibitor (NSC-87877) showed modestly decreased [H+], slightly elevated acute ROS, and no exocytosis (Figure 6D-F). Together, the data show that SHP-1 deficiency or inhibitors of SHP-1 activity are not sufficient to fully recapitulate the LEL dysfunction of MRL/*lpr* mice.

Phosphorylation of SHP-1 at Y^564^ (pSHP-1) is necessary for phosphatase activity (41) Stimulation of B6 and MRL/*lpr* BMMφs with IgG-IC (15min – 5hr) induced comparable pSHP-1^Y564^ at all time point

(Supplemental Figure 8A). Since FcγRI constitutively resides within lipid rafts (42), SHP-1 must localize to lipid rafts to dephosphorylate the FcγRI ITAM (43). We used confocal microscopy to assess colocalization of pSHP-1^Y564^ with cholera toxin (CTx) stained membrane lipid rafts in B6 and MRL/*lpr* BMMφ following IgG-IC stimulation. We found comparable co-localization of pSHP-1^Y564^ with lipid rafts in B6 and MRL/*lpr* BMMφ (Supplemental Figure 8B and C). Thus, reduced SHP-1 phosphorylation, or intracellular pSHP-1^Y564^ mislocalization, does not impair LEL function in MRL/*lpr* BMMφ. Reduced SHIP-1 and SHP-1 activity impairs LEL function

To assess whether deficiency in both SHP-1 and SHIP-1 recapitulate LEL dysfunction in MRL/*lpr* mice, we treated B6.SHIP-1^-/-^ BMMφs with the SHP-1 inhibitor NSC87877. After stimulation with IgG-ICs, (30 min) B6.SHIP-1^-/-^ BMMφs treated with SHP-1 inhibitor showed 4.5x-fold (p=0.05) lower [H+], while MRL/*lpr* BMMφ show 10.3-fold (p=0.0001) lower compared to B6 t_30_ (Figure 6G). Further, B6.SHIP-1^-/-^BMMφs treated with SHP-1 inhibitor showed 2.7-fold increase in ROS (15 min, p=0.04), while MRL/*lpr* BMMφs showed a 2.1-fold (p=0.03) increase compared to untreated cells (Figure 6H). Sustained ROS (2hrs) in B6.SHIP-1^-/-^+SHP-1 inhibitor was also elevated (2.2-fold increase, p=0.065) while MRL/*lpr* levels were 2.0-fold (p=0.04). Although B6.SHIP-1^-/-^+SHP1/2 inhibitor diminished acidification and heightened ROS, it did not induce exocytosis of undegraded IgG-ICs (Figure 6I).

To corroborate these findings, we treated B6.SHP-1^-/-^ BMMφs with SHIP-1 inhibitor (3AC). Compared to B6, IgG-IC stimulation of B6.SHP-1^-/-^ BMMφs with SHIP-1 inhibitor showed a 9.1-fold lower [H+] at 30 min (p=0.003), while MRL/*lpr* BMMφ was 10.3-fold increased (p=0.0001; Figure 6G). Additionally, B6.SHP-1^-/-^ BMMφs treated with SHIP-1 inhibitor for 15 min showed 8.4-fold heightened ROS (p=0.002) while MRL/*lpr* showed 2.1-fold (p=0.03; Figure 6H). Levels of sustained ROS (2 hrs) were 10.7-fold increased (p=0.001), while MRL/*lpr* was 2.0-fold increased (p=0.04). The dual deficiency of SHP-1 and SHIP-1 induced a 2.6-fold increase in the exocytosis of undegraded IgG-ICs (p=0.02; Figure 6I). Thus, LEL dysfunction in BMMφs from B6.SHP-1^-/-^+SHIP-1 inhibitor recapitulated d MRL/*lpr*, while B6.SHIP-1^-/-^+SHP-1 inhibitor was less effective. One possibility is that although heightened ROS is necessary, it is not sufficient to drive LEL dysfunction, while diminished acidification plays a more important role. Collectively, the data show that defects in SHP-1 and SHIP-1 contribute to LEL dysfunction in MRL/*lpr* mice suggesting these phosphatases play direct or indirect roles in disabling LEL function in lupus. Whether other phosphatases in the PI3k/Akt pathway (PTEN, Akt phosphatases (PP2A, PHLPP1/2)) also play similar roles is unclear.

## Discussion

Our murine studies show that activation of PI3K, in part through chronic FcγRs signaling results from, and leads to, diminished LEL acidification creating a feedforward loop (13–15). In human disease, non-acidic LELs and increased circulating and membrane-bound IgG-ICs are evident during active disease, while acidic LELs and low levels of circulating and membrane-bound IgG-ICs prevail during inactive disease. This is consistent with the idea that LEL dysfunction could drive active versus inactive disease. In further support of that idea, we find diminished [H+] and hydrolase activity are evident in patients with moderate and highly active SLE (Figure 3), yet the accumulation of surface nucleosome and increased CIC are predominately seen in highly active disease (Figure 4). These data suggest that non-acidic LELs decrease the degradation of IgG-ICs, which in turn promotes exocytosis and heightens CIC and the accumulation of undegraded IgG-ICs on the plasma membrane (9). Thus, LEL dysfunction is unlikely the ‘cause of lupus’; rather, an induced defect that perpetuates disease (Figure 5C-D). This is supported by GWAS and twin studies demonstrating genetic underpinnings in SLE (1, 44). Clinically, LEL dysfunction is associated with increasing SLE disease activity (Figure 3G-I, 4I-M) and is coincidental with, or may precede, clinical manifestations of SLEDAI rash, arthritis, and nephritis (Table S4). Non-acidic LELs are evident in 67% of SLE patients (65% Mo, 57% DCs), raising the possibility that restoring LEL function could be a therapeutic target that sustains inactive disease.

Murine studies showed that diminished LEL acidification is induced through PI3k activation (Figure 5C and D). This suggests that heritability of LEL dysfunction could originate from genes conferring indirect effects that disrupt cellular signaling networks required to regulate immune responses (45). The data also suggest that FcγRI signaling, induced by newly formed IgG-ICs, or those that have accumulated on the cell surface, is not properly terminated and chronically activate the PI3k/Akt/mTORC pathway (Figure 5). Chronic activation of FcγRI/PI3K/Akt/mTOR that heightens ROS and reduces acidification is also sustained by disrupted regulation by SHP-1 and SHIP-1, phosphatases that normally attenuate FcγRI signal transduction (Figure 6). Genetic polymorphisms could also disrupt LEL function. Transient ROS production by nicotinamide adenine dinucleotide phosphate (NADPH) oxidase is important for LEL maturation and acidification (46), deficiency in NOX2 exacerbates murine SLE (47), and genetic risk variants of NCF that lower ROS promote lupus in non-autoimmune mice (48, 49). In contrast, heightened and/or prolonged ROS impairs the activity of lysosomal hydrolases (50), diminishes activity of protein tyrosine phosphatases through oxidation at the catalytic site of protein, (51), and diminishes phagosomal acidification by disrupting V-ATPase assembly (52). Thus, ROS is tightly regulated to allow transient increases that are appropriately terminated. In summary, genetic and induced events can disrupt LEL function in SLE, with our human and murine data revealing a potential mechanism underlying an induced defect.

In non-autoimmune B6 mice, deficiency in both SHIP-1 and SHP-1 is sufficient to reduce LEL acidification, heighten ROS, and promote exocytosis of undegraded IgG-ICs (Figure 6G-I). SHP-1 dephosphorylates Y^58^ within the ITAM of FcRγ and multiple signaling effectors that regulate FcRγ signaling, including src family kinases pSyk, p62Dok, and pVav. One possible mechanism by which SHP-1 might contribute to LEL dysfunction in MRL/*lpr* mice is through reduced phosphatase activity due to high ROS that chronically oxidizing the SHP-1 catalytic site thereby limiting dephosphorylation of FcγRI ITAM tyrosines and/or other receptor proximal signaling effectors.

SHIP-1 also plays a crucial role in LEL dysfunction in MRL/*lpr* mice (Figure 5 E-J, 6A-C). SHIP-1 regulates the levels of PIP3, which act as a docking site for kinases (53) regulating FcγRI activation at the level of PI3k activation (Figure 5E). In MRL/*lpr* mice, diminished phosphorylation of SHIP-1 could prevent FcγRIIB/SHIP-1co-localization with FcγRI-containing lipid rafts. This would sustain PIP3 levels and induce LEL dysfunction. Exclusion of FcγRIIb/SHIP-1 from lipid rafts might occur through mutations within the FcγRIIb transmembrane region (54, 55), or through altered composition of glycosphingolipids, as in CD4+ T cells lipid rafts from SLE patients (56). Last, we did not see a decrease in SHIP-1 total protein, thus, do not think microRNA regulation of SHIP-1 is involved.

How does LEL dysfunction occur in lymphocytes that don’t typically express activating FcγRs? LEL dysfunction in B cells could involve FcγRIIb and surface receptors that bind apoptotic debris like autoreactive BCR, or receptors that bind opsonins such as complement, oxLDL, Gas6, or milk fat globule epidermal growth factor 8. In B cells, late endosomes are the preferred site for degrading endocytic cargo (57) and accumulation of undegraded nucleic acid drives TLR7 pathologies in SLE (58, 59). T cells also do not typically express activating FcγR; however, autophagy defects (60) affect T cell metabolism and mitochondrial function (61, 62). Autophagy requires functional LELs to degrade intracellular waste. A therapeutic phospho-peptide derived from the spliceosomal U1-70K protein reduces chaperone-mediated autophagy and limits T cell activation and plasma cell formation in murine and human SLE (63). In summary, LEL defects are evident in both murine and human lupus and inducible via chronic FcγRI/PI3K/Akt/mTOR signaling from newly formed and/or exocytosed IgG-ICs. The sustained activation of PI3K pathway is (or could) in part be due to disrupted SHIP1 and SHP1 function. Thus, a feedforward loop, created when elevated IgG-ICs chronically activate FcγR/PI3k signaling, reduces LEL acidification, heightens ROS and sustains IgG-IC levels through exocytosis of undegraded LEL cargo, setting in motion a sustained cycle of LEL dysfunction and inflammation.

<u>Study limitations</u> include the relatively low enrollment of Hispanic and Asian patients, the cross-sectional nature of the study. We chose hybrid SELENA-SLEDAI as the disease activity marker because it is a validated, reliable, objective SLE disease activity measure that is easy to score and not confounded by fibromyalgia. The well accepted disadvantage of SLEDAI is insensitivity to small change, and inability to capture severity within each category. The strengths of this study are that our demographics reflect the overall SLE population with a predominance of Black females, we enrolled SLE and RA patients from two sites, and SLE patients were grouped based on disease activity (inactive, moderately active, highly active).

## Methods

### Sex and biological variable

For human studies, sex was not considered as biological variable. For studies with animals, both genders were used since both male and female MRL/*lpr* mice develop spontaneous lupus.

### Study Design and Demographics

Peripheral blood hematopoietic cells were analyzed at UNC. At the time of visit, disease activity was scored using hybrid SELENA-SLEDAI (64). Disease activity and demographic data were entered into UNC or Duke REDCap databases and provided in Supplemental Tables 2, 6, and 9. SELENA-SLEDAI or DAS 28 scores were not available until after laboratory data were analyzed, and the clinical investigators were blinded to the results of the study until final data analysis were completed. No duplicate patients were used in the study.

### Patient Enrollment Criteria

Consenting patients who met the 1997 ACR or 2012 SLICC criteria for SLE (65) were enrolled. Exclusion criteria (prospective) included under 18 years, chronic infection, active malignancy, inability to provide informed consent, or receiving rituximab within 6 months, and IVIG, belimumab or a biologic medication for rheumatic disease (including anti-TNF therapy) within 3 months. The inclusion criteria for rheumatoid arthritis were that patients 1.) met ACR RA criteria, 2.) showed seropositivity for RF or CCP, 3.) showed activity as measured by DAS28 ESR ≥3.2 or DAS28 CRP ≥2.9, 4.) had at least 1 joint having synovitis, and 5.) were not on biologics or small molecule therapy. Healthy controls were enrolled through the UNC Platelet Donation Center or Duke OB-GYN clinic and selected as those without family history of autoimmune disorders or current symptoms of infection.

### Animal Models

We obtained C57BL/6 (B6), MRL/MpJ-*Tnfrs6^lpr^*/J (MRL/*lpr*; 000485), MRL/MpJ (000486) and B6.MRL-Fas^lpr^/J (B6.*lpr*; 000482) mice from Jackson Laboratories, C57BL/6-*Gt(ROSA)26Sor^tm9(Cre/ESR1)Arte^* (B6.Rosa26-CreERT2, 10471) from Taconic Biosciences, and B6.SHP-1^fl/fl^ mice from Paul Love (NIH, Bethesda MD). FcγRI^-/-^/MRL/*lpr* were previously described (13). When mice reached disease endpoint described in IACUC protocol, they were euthanized following IACUC guidelines.

### Reagents and Antibodies

The list is provided in Supplemental Methods.

### IgG-Immune Complex (IgG-ICs)

IgG-ICs were made from apoptotic blebs obtained by irradiating B6 thymocytes (7-9 weeks mice), or human thymoma cells CCRF-CEM (180×10^6^, ATCC #CCL-119) at 600 rads in 10ml of PBS + 2% FBS. After incubation (16-18 hrs), supernatants containing apoptotic debris were collected from centrifuged (350 xg) cultures. Depending on sample number, a given volume of the apoptotic debris was incubated with 12 μg/ml of PL2-3 (IgG2a, (13)) for 1 hr (mouse study), or 12 μg/ml 33H11 (IgG1; (66)) for 2 hrs (human study) at room temperature. IgG-ICs were pelleted by centrifuging at 160,000 x*g* for 45 min at 4°C, and resuspended in <u>complete media</u> (10% FBS, 1 mM sodium pyruvate, 50 μg/ml gentamicin, 100 U/ml penicillin, 100 μg/ml Streptomycin, 2 mM L-glutamine, and 5×10^-5^ M β-mercaptoethanol in phenol-red free RPMI for human samples or phenol-red free DMEM for mouse samples) at 25% of the volume that was used in binding PL2-3/33H11. To stimulate cells, we used 25 or 30 μl of IgG-ICs/0.25×10^6^ cells.

### Flow Cytometry

RBC lysed blood cells from human samples and mouse splenocytes, and BMMφs were stained for surface bound and intracellular targets using standard methods for flow cytometry (Supplemental Methods).

To inhibit PI3K p110 subunit isoforms, BMMφs were treated with 100nM of PIK-75 (inhibit p110α with 13-fold higher specificity than p110γ), TGX-221 (inhibit p110β with 20-fold higher specificity than p110δ), and IC-87114 (inhibit p110δ with 58-fold higher specificity than p110γ), 2hrs prior to IgG-IC treatment. To inhibit SHIP1, we used 3AC (50nM) on day 5 (48hrs prior to IgG-IC treatment) and for SHP1, we used NSC87877 (10μM) at 3hrs prior to IgG-ICs treatment. To generate SHP1 deficient BMMφs, B6.SHP-1^fl/fl^ x B6.Rosa26-CreERT2 were treated with tamoxifen (2mg in 100μl corn oil, i.p.) for 4 consecutive days, then at 17days post 1^st^ injection of tamoxifen, bone marrow cells were collected and cultured as described above. SHIP1 deficient BMMφs were generated from the bones of B6.SHIP1^-/-^ mice provided by Gerald Krystal (British Columbia Cancer Agency, Vancouver, BC, Canada). We obtained spleens and bones of NZM2410 mice from Melissa Cunningham (Medical University of South Carolina, Charleston), and of B6.Sle123 mice from Laurence Morel (Univ of Texas, San Antonio).

#### a. Late endosome/lysosome pH

Unfractionated RBC-lysed human blood cells or mouse splenocytes/BMMφs (0.25×10^6^) were stained for cell markers (Supplemental Methods). Cells were stimulated with IgG-ICs (25 or 30 μl of IgG-ICs/0.25×10^6^cells), for 30 min at 37°C, then LEL pH was measured. Samples for 60min were washed and resuspended in 200μl of warmed complete media, then continued incubating at 37°C for 30 min. To establish time 0 (t_0_), the vAPTase was inhibited using Concanamycin A (human: 20 ng/ml, mouse: 2 ng/ml). To measure pH changes, LysoSensor 160-DND (1μl/sample) was added 15 min prior to flow cytometry. LysoSensor was excited with a UV laser (355 nm) and relative pH was calculated by ratioing the MFI from two emission channels (450/20 nm, 585/42 nm). Linear regression from standard curve of each cell type was used to calculate absolute pH, followed by conversion to hydrogen ion concentration ([H+]) based on pH = -log_10_[H+]. The standard curve was generated using intracellular pH calibration buffer.

#### b. Late endosome/lysosome hydrolase activity

Late endosomal/lysosomal hydrolase activity was measured using hydrolase activity kit with a modified protocol. In brief, cells were treated with IgG-ICs (30 μl of IgG-ICs/0.25×10^6^ cells) and the “self-quenched substrate” (15 μl/1 ml) for 60 min at 37°C. After washing, cells were left at room temperature for 45 min, then analyzed by flow cytometry. The MFI at 488 nm reflected hydrolase activity. Relative hydrolase activity was calculated by subtracting MFI of fluorescent minus one (FMO) for FITC, then normalized to Concanamycin A treated sample (t_0_).

#### c. ROS

Intracellular ROS levels were measured using CellROX Deep Red kit with modified protocol. In brief, while BMMφs were treated with IgG-ICs (25 μl of IgG-ICs/0.25×10^6^ cells) for 15 and 120 min, CellROX (1μM) was added with antibodies for cell markers 15 min before the end of IgG-IC treatment. Cells were stained with LIVE/DEAD kit, then fixed in 2% PFA /FACS buffer. Samples without IgG-IC stimulation are t0 samples. ROS levels were calculated by subtracting MFI of FMO from MFI of CellROX stained samples.

#### d. Exocytosis of fluorochrome labeled IgG-ICs to plasma membrane

IgG-ICs were made with apoptotic debris and AlexaFluoro488 (AF488) labeled PL2-3. BMMφs were pre-loaded with AF488-labeled IgG-ICs for 30 min at 4°C and unbound IgG-ICs were removed by washing. After incubation at 37°C for 24 and 72hrs (± inhibitors for some experiments), half of the samples were treated with anti-AF488 antibody to quench surface bound AF488 fluorescence and the other half were left unquenched. To set t0, cells pre-loaded with AF488 labeled IgG-ICs were processed the same way with anti-AF488 without further incubation. All samples were then fixed and analyzed by flow cytometry. MFI from quenched samples indicate fluorescence from internalized IgG-ICs and MFI of unquenched samples depict total fluorescence from both surface bound and internalized IgG-ICs. Levels of exocytosed IgG-ICs were calculated by subtraction (internalized and surface fluorescence minus internalized fluorescence). MFI of exocytosed IgG-ICs were normalized to individual t0.

#### f. Circulating immune complex (CIC)

The concentration of CICs were quantified by measuring the loss of FcγRIIA when HC neutrophils were incubated with patient (or HC) plasma (67). We separated plasma on the day of blood collection, aliquoted, and stored at -80°C until analysis. Neutrophils were isolated from HC blood using Polymorphprep. After RBC lysis with ACK buffer, 0.2×10^6^ cells/180 μl of phenol red-free RPMI were plated into a 96-well U-bottom plates and 20 μl of patient plasma was added to each well (final 10%). Plates were incubated (37°C, 90 min) to allow for internalization of the plasma immune complexes. To stop internalization, cells were washed with cold FACs buffer containing 0.2% of NaN_3_, then stained with fixable Live/Dead. To measure surface expression of FcγRIIA, cells were stained with fluorochrome conjugated IV.3 (anti-FcγRIIA) and antibodies for cell markes, then fixed. We measured the MFI of IV.3 staining on CD45^pos^CD14^neg^HLA-DR^neg^CD16^pos^CD11b^pos^CD15^pos^ neutrophils. The concentration of plasma CIC was calculated using a standard curve generated by treating cells with various concentrations of heat-aggregated IgG-ICs (human 33H11 heated 1hr at 63°C).

Samples were acquired on Thermo Fisher Attune NxT or Cytek Aurora (5 lasers). Lysosome pH and hydrolase activity data for the human study were obtained on Becton Dickinson LSR Fortessa, and for the murine study on Becton Dickinson LSRII. Data acquired from Cytek Aurora were unmixed using Spectroflo. All flow cytometry data were analyzed with FlowJo and graphed using GraphPad Prism.

### Confocal microscopy

We used confocal microscopy to measure colocalization of cholera toxin-stained membrane lipid rafts with intracellular pSHIP1^Y1022^ or pSHP1^Y564^ (Supplemental methods). Quantitative data of colocalization was obtained by calculating Mander’s coefficient of colocalization (colocalized pixels/total fluorescent pixels within region of interest).

### Statistical Analysis

Murine data were analyzed using the non-parametric 2-way ANOVA, Kruskal-Wallis, or Mann-Whitney test with original False Discovery Rate (FDR) method (Benjamini-Hochberg, [BH]) for multiple comparisons (GraphPad Prism v9.5.1). For human data, we used Shapiro-Wilk test, visual inspection of Q-Q plots, and Levene’s test to assess homogeneity of variance to assess normality of data. We used Dunn (1964) Kruskal-Wallis nonparametric ANOVA adjusted for FDR (BH), followed by pairwise comparisons to assess the differences between four disease groups (HC, SLE patients with hybrid SELENA-SLEDAI (SLEDAI) of ≤5, 6-11, or ≥12). Cochran-Armitage test was used to identify trends between the proportion of patients in each disease activity group and parameters associated with LEL dysfunction ([H+], hydrolase, surface nucleosome, CIC). The p-values were adjusted for multiple comparisons (BH method). Fold change was used in the text for ease of comparison between SLEDAI groups; however, fold change was not used in calculating p-values. The statistical methods for calculating p-values were provided in each figure legend and were done between all groups or experimental conditions, but only two-sided p-values <0.05 were considered statistically significant. Associations between SLEDAI groups and the presence of diminished [H+] and hydrolase activity, or elevated surface nucleosome and circulating immune complexes were defined by cut-offs: <u>non-acidic</u>: patients with non-acidic lysosomes have [H+] lower than the cut-off. The cut-off for each cell type was established at 1.8-fold above the mean [H+] of the HC, <u>low hydrolase</u> patients have levels below a cut-off that was established at 1.7-fold above the mean hydrolase of HC for each cell type, <u>high surface nucleosome</u>: patients with high surface nucleosome have cell surface nucleosome levels above the cut-off. The cut-offs for each cell type were established at 1.8-fold above the mean surface nucleosome level of the HC, <u>high CIC</u>: patients with high CIC show plasma CIC levels higher than the cut-off. The cut-off was established at 1.5-fold above the mean CIC level of the HC. These are also described in the footer of Table S4. Clinical categories were described with mean ±SD for continuous variables, and N (%) for binary categorical variables. The study was not powered for comparisons between clinical categories, so these comparisons were not made to avoid inflating type II error.

### Study approval

Primary Institutional Review Board oversight was the responsibility of the UNC Office of Human Research Ethics with approval of the consent form and protocols for the study by Duke University Health System - Institutional Review Board for Clinical Investigations. Participants were enrolled under approved UNC IRBs (12-2097, 18-3193, 19-1690), and Duke IRBs (Pro00008875, Pro0000775, Pro00094645, Pro00080944). During routine clinic visits, patients providing approval of written consenting form were enrolled. This study was approved under UNC Institutional Animal Care (IACUC 21-136, 24-092).

## Data availability

All experimental data are shown in the main text or the supplementary materials, and are provided in the Supporting Data Values file.

## Author Contribution

SAK, JLR, BJV designed the study. SAK, AJM conducted experiments, acquired and analyzed data. JLR (Duke), SZS (UNC) identified patients. KG (Duke), SSB, ACT (UNC) consented patients. KG, SSB, ACT, XB procured samples. KS, RES, MM, MEBC, JLR (Duke) and SZS (UNC) provided clinical commitment. BJV, JLR oversaw the project. LA performed statistical analysis. SAK, BJV wrote the manuscript. We determined the order of authorship based on the levels of contribution to the project.

## Funding Support

National Institutes of Health (NIH) 1R01AI132421-01A1 (BJV, JLR, MEBC) NIH P30CA016086, 1UM2AI30836-01, 5P30AI05041 (Flow Cytometry Core)

NIH Center for Clinical Research P30AR072580 (Thurston Arthritis Center-Statistical Core)

NIH Center for Advancing Translational Sciences Clinical and Translational Science UL1TR002489 (REDCap data management)

Lupus Research Alliance Novel Research Grant (BJV, JLR) Department of Defense Lupus Research Program (BJV, SZS)

North Carolina Biotechnology Center – Biotechnology Innovation Grant (BJV)

North Carolina Biotechnology Center IDG-1025 (Flow Cytometry Core)

Rosemarie K Witter Foundation Inc – gift (BJV)

## Acknowledgements

General:

We thank Amanda Nelson for critically reviewing this manuscript, the Duke and UNC Lupus Registries and the SLE patients who generously provide blood for this study, the UNC Platelet Donation Center and staff for providing healthy control blood, UNC Microscopy Services Laboratory, Thomas H Winkler for providing 33H11 (anti-DNA) hybridoma, Quan Li Zhen for confirming specificity of 33H11, Christian Lood for advice establishing the flow-based CIC assay, and Edna Scarlett and Julie Walker for managing the Duke and UNC IRBs. Graphical abstract was created in BioRender.

## Conflict of interest statement

SAK, AJM, LA, KG, SSB, ACT, XB, KS, MM, SZS declares no conflict of interest. RES reports grants/contracts from Rheumatology Research Foundation and Lupus Research Alliance and consulting fees from Childhood Arthritis and Rheumatology Research Association. MEBC reports grants from GSK and UCB. JLR declares grants or contracts from Exagen, Immunovant, and the Department of Defense. BJV reports a pending patent application, incorporation of Autoimmune Therapeutics, and monetary donations from the Rosemarie K Witter Foundation.

## Abbreviations

aNAV: activated naïve B cells
APC: antigen presenting cell
BCR: B cell receptor
BH: Benjamini and Hochberg method for
FDR: False Discovery Rate
BMMφ: bone marrow-derived Mφ
CIC: circulating immune complexes
DCs: dendritic cells disease activity group – inactive; SLEDAI ≤5, moderately active; SLEDAI 6-11 highly active; SLEDAI ≥12)
DMARD: disease-modifying anti-rheumatic drugs
DN2: double negative-2 B cells
dsDNA: double stranded DNA
endosomes/lysosomes: endosomes and lysosomes
FcgR: Fc gamma receptor
[H+]: hydrogen ion concentration
HCQ: hydroxychloroquine
LELs: late endosomes and lysosomes
Mφ: macrophages
Mo: monocytes
SLE: systemic lupus erythematosus
SLEDAI: SLE Disease Activity Index from hybrid SELENA-SLEDAI
(t0): time 0 min
QTLs: quantitative trait loci

