## Supplemental data and tables for "Reduced late endosome/lysosome function promotes SLE through chronic PI3k activity and SHP-1/SHIP-1 defects"

### Supplemental Materials

### Supplemental Methods

#### *Reagents and Antibodies*

| Reagent | Company | Catalog # |
|---|---|---|
| Concanamycin A | Sigma | C9705 |
| Intracellular pH Calibration Buffer kit | Thermo Fisher Scientific | P35379 |
| Live/Dead Fixable Near IR (780) Viability Kit | Thermo Fisher Scientific | L34994 |
| LysoSensor Yellow/Blue DND-160 | Thermo Fisher Scientific | L7545 |
| Hydrolase Activity Assay Kit | Abcam | Ab234622 |
| Phenol-red free High Glucose DMEM | Gibco | 21063029 |
| Phenol-red free RPMI | Gibco | 11835030 |
| Polymorphprep | Fisher scientific | NC0863559 |
| Tamoxifen | Selleckchem | S1238 |
| 3AC | Echelon Biosciences | B-0341 |
| NSC87877 | Selleckchem | S8182 |
| PIK-75 HCl | Selleckchem | S1205 |
| TGX-221 | Selleckchem | S1169 |
| IC-87114 | Selleckchem | S1268 |
| CellROX Deep Red Flow Cytometry Assay kit | Thermo Fisher Scientific | C10491 |
| Cholera Toxin Subunit B AlexaFluor 594 conjugated | Thermo Fisher Scientific | C34777 |

| Antibody Target | Fluorochrome | Company | Catalog # | RRID |
|---|---|---|---|---|
| CD19 | PE | Biolegend | 392506 | AB_2750097 |
| CD3 | BV650 | Biolegend | 300468 | AB_2629574 |
| CD3 | PE-Dazzle 594 | Biolegend | 300336 | AB_2632652 |
| CD14 | AF 488 | Biolegend | 301811 | AB_493159 |
| CD14 | APC-Cy7 | Biolegend | 301820 | AB_493695 |
| CD16 | PECy7 | Biolegend | 302016 | AB_314216 |
| CD11c | PE-CF 594 | BD Biosciences | 562393 | AB_11153662 |
| CD11c | APC | BD Biosciences | 559877 | AB_398680 |
| HLA-DR | BV421 | Biolegend | 307636 | AB_2561831 |
| HLA-DR | AF 488 | Biolegend | 307620 | AB_493175 |
| HLA-DR | BUV395 | BD Biosciences | 564040 | AB_2738558 |
| CD66b | PerCP-Cy5.5 | Biolegend | 396914 | AB_2820061 |
| CD56 | PerCP-Cy5.5 | Biolegend | 392420 | AB_2734444 |
| CD24 | Superbright 600 | Thermo Fisher Scientific | 63-0247-42 | AB_2717025 |
| CD21 | PECy7 | BD Biosciences | 561374 | AB_10681717 |
| IgD | APC-H7 | Biolegend | 348218 | AB_11203722 |
| CXCR5 | AF 488 | R and D Systems | FAB190G-100UG | AB_3647470 |
| CD38 | PerCP-Cy5.5 | BD Biosciences | 551400 | AB_394184 |
| CD27 | BV510 | BD Biosciences | 563092 | AB_2313577 |
| CD64 | AF 647 | Biolegend | 305012 | AB_528867 |
| CD16 | AF 647 | Biolegend | 302020 | AB_492976 |
| CD32A | AF 647 | Novus | NBP2-47830AF647 | AB_3312796 |
| CD32B/C | APC | Biolegend | 398304 | AB_2860997 |
| CD15 | BV605 | Biolegend | 323031 | AB_2562131 |
| CD45 | PE | Biolegend | 304058 | AB_2564156 |
| CD32A (clone IV.3) | Unconjugated | Stemcell Technologies | 60012 | AB_2925215 |
| Ly6G | BUV563 | Thermo Fisher Scientific | 365-5931-82 | AB_2925400 |
| Ly6G | AF 488 | Biolegend | 127625 | AB_2561339 |
| CD11b | Spark Plus UV385 | Biolegend | 101240 | AB_3097646 |
| CD11b | APC | BIolegend | 101212 | AB_312795 |
| F4/80 | eFluor 450 | Thermo Fisher Scientific | 48-4801-82 | AB_1548747 |
| CD19 | BUV737 | Thermo Fisher Scientific | 367-0193-82 | AB_2895945 |
| CD3 | BV785 | Biolegend | 100232 | AB_2562554 |
| SHP-1 | - | Cell Signaling Technology | 3759S | AB_2173694 |
| pSHP-1 (Tyr564) | - | Cell Signaling Technology | 8849S | AB_11141050 |
| SHIP-1 | - | Cell Signaling Technology | 2728S | AB_2126244 |
| pSHIP-1 (Tyr1022) | - | Cell Signaling Technology | 3941S | AB_2296062 |
| Anti-Rabbit IgG | AF 647 | Thermo Fisher Scientific | A-21244 | AB_2535812 |
| PIP3 | Fluorescein | Echelon Biosciences | Z-G345 | AB_427218 |
| PI(3,4)P2 | - | Echelon Biosciences | Z-P034 | Unavailable |

#### *PCR*

B6.SHP-1$^{fl/fl}$ x B6.Rosa26-CreERT2 were genotyped for CreERT2 using primers for Cre negative forward 5' CTG GCT TCT GAG GAC CG 3', Cre negative reverse 5' CCG AAA ATC TGT GGG AAG TC 3', Cre positive forward 5' CGT GAT CTG CAA CTC CAG TC 3', Cre positive reverse 5' AGG CAA ATT TTG GTG TAC GG 3'.

#### *Cell markers for flow cytometry*

**Mouse cell markers**

| Cell Types | Markers |
|---|---|
| B cells | CD3$^{neg}$CD19$^{pos}$ |
| T cells | CD19$^{neg}$CD3$^{pos}$ |
| CD11b myeloid cells | CD19$^{neg}$CD3$^{neg}$CD11b$^{pos}$ |
| CD11c+ DCs | CD19$^{neg}$CD3$^{neg}$CD11c$^{high}$ |
| Neutrophils | CD19$^{neg}$CD3$^{neg}$CD11b$^{pos}$Ly6G$^{pos}$ |
| BMMφs | CD11b$^{pos}$F4/80$^{pos}$ |

neg = negative, pos = positive

**Human cell markers**

| Cell Types | Markers |
|---|---|
| Monocyte | CD19$^{neg}$CD3$^{neg}$CD66b$^{neg}$CD56$^{neg}$CD14$^{pos}$HLA-DR$^{pos}$ |
| DCs | CD19$^{neg}$CD3$^{neg}$CD66b$^{neg}$CD56$^{neg}$CD14$^{neg}$CD11c$^{pos}$ |
| B cells | CD66b$^{neg}$CD56$^{neg}$CD3$^{neg}$CD19$^{pos}$ |
| T cells | CD66b$^{neg}$CD56$^{neg}$CD19$^{neg}$CD3$^{pos}$ |
| rNAV | CD19$^{pos}$CD27$^{neg}$IgD$^{pos}$CD24$^{neg}$CD38$^{neg}$CD21$^{pos}$CD11c$^{neg}$ |
| aNAV | CD19$^{pos}$CD27$^{neg}$IgD$^{pos}$CD24$^{neg}$CD38$^{neg}$CD21$^{neg}$CD11c$^{pos}$ |
| DN1 | CD19$^{pos}$CD27$^{neg}$IgD$^{neg}$CD21$^{pos}$CD11c$^{neg}$ |
| DN2 | CD19$^{pos}$CD27$^{neg}$IgD$^{neg}$CD21$^{neg}$CD11c$^{pos}$ |

neg = negative; pos = positive

#### *Flow cytometry*

Blood cells were collected after RBC lysis with ACK buffer (0.15 M $NH_4Cl$, 1 M $KHCO_3$, 0.1 mM $Na_2EDTA$, pH 7.2-7.4), or for mouse splenocytes after RBC lysis with $NH_4Cl_2$ (0.154 M, pH 7.2). Cells were washed twice with PBS, filtered through 40μm strainer, and resuspended in complete media. BMMφs from 8-10 wk old mice were derived as previously described {Monteith, 2016 #4}. In short, bone marrow cells from tibias and femurs were grown in complete media with 3-5% L-cell supernatant for 7 days with media replenishment on day 4. Day 7 BMMφs (95-98% CD11b$^{pos}$F4/80$^{pos}$) were collected and rested for 2hrs at

37°C before treatment. Human unfractionated blood cells ($1x10^6$) or mouse splenocytes/BMMφs ($1x10^6$) were blocked with 30% human AB serum or 200 μg/ml of 2.4G2 (anti-FcγRIIB/III) for 15 min on ice. Cells were washed in FACS buffer (1x PBS with 2% FBS, 0.2% $NaN_3$) and stained for surface bound nucleosome (PL2-3, mouse IgG2a, conjugated with Alexafluor 647), dsDNA (33H11, human IgG1, conjugated with Alexafluor 647), FcγRI, FcγRIIa, FcγRIIb, and FcγRIII in the presence of antibodies to various cell markers, then fixed with 2% paraformaldehyde (PFA) in FACS buffer. For intracellular levels of $pSHIP1^{Y1022}$, SHIP1, $pSHP1^{Y564}$, SHP1, $PIP_3$, and $PI_{(3,4)}P_2$, cells were stained for cell markers, fixed in 3% PFA, permeabilized with 0.2% saponin in FACS buffer, then stained for intracellular antigens. All samples were stained with Live/Dead Fixable Viability Kit. Data are presented as mean fluorescence intensity (MFI) of sample/MFI of isotype antibody.

##### *Confocal microscopy*

BMMφs were treated with fluorochrome-labeled IgG-ICs (30 min, 37°C), then stained (10 min, 4°C) with cholera toxin B subunit conjugated with AF594 (CTx) to identify membrane lipid rafts. Cells were fixed in 2% PFA in PBS, then permeabilized with 0.2% saponin in FACS buffer. Cells were blocked with 5% rat serum in FACS buffer, then stained for intracellular $pSHIP1^{Y1022}$ or $pSHP1^{Y564}$. Cells were resuspended in FluorSave and loaded onto coverslips for imaging. Images were acquired using Zeiss 710 confocal microscope with a 63x 1.4 N.A. (oil) PLAN APO lens and Zeiss Zen software. Cells were imaged from many fields randomly selected across the coverslip. Cell clumps or poorly stained cells were excluded. Images were processed using Image J. Colocalization of $pSHIP1^{Y1022}$ or $pSHP1^{Y564}$ with CTx positive lipid rafts were quantified using the Mander's coefficient of colocalization (colocalized pixels/total fluorescent pixels within region of interest).

#### Supplemental Figures

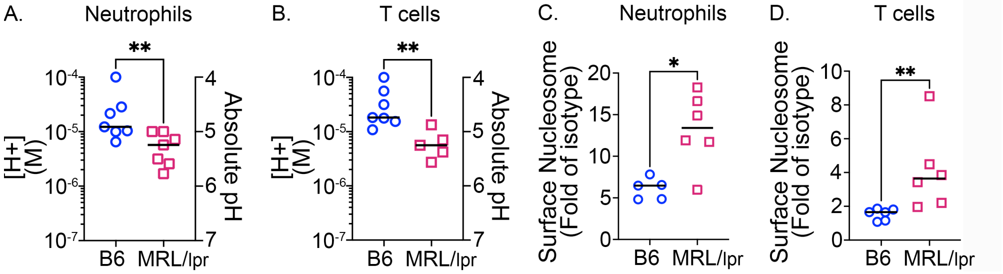


**Supplemental Figure 1. T cells and neutrophils from MRL/*lpr* mice show diminished [H+] and elevated surface nucleosome.** LEL [H+] and the levels of surface nucleosome were analyzed on splenic neutrophils ($CD45^+CD19^-CD3^-CD11b^+Ly6G^+$, **A, C**) and T cells ($CD45^+CD19^-CD3^+$, **B, D**). Splenocytes were incubated at 37°C in the presence (for neutrophils) or absence (for T cells) of IgG-ICs (30μl IgG-ICs/$0.25 \times 10^6$ cells) for 30min and LEL pH was measured by flow cytometry. Absolute pH was calculated using a standard curve, then converted to [H+] using pH= $-\log_{10}$[H+] **(A, B)**. N = 5-7, 3 experiments. Statistical analysis used Mann-Whitney test. *$p<0.05$, **$p<0.01$, ***$p<0.001$. Bar = median.

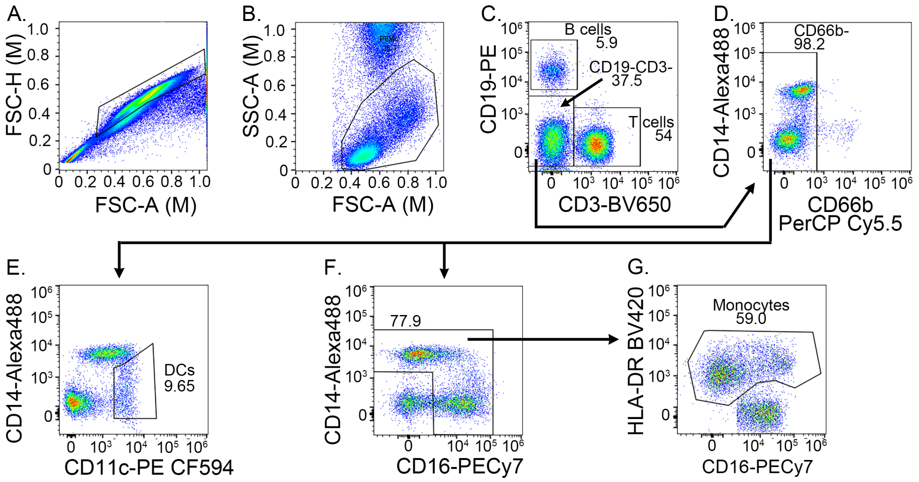


**Supplemental Figure 2. Flow cytometry gating of human blood hematopoietic cells.** Unfractionated blood cells were stained with antibodies specific for CD19, CD3, CD14, CD66b, CD11c, CD16, and HLA-DR. Singlets **(A),** were further gated for lymphocytes plus monocytes (lymph-mono gate) **(B)**, then subsequently gated for B and T cells using CD19 and CD3 **(C)**. CD19$^{neg}$CD3$^{neg}$ cells were further gated for CD66b to exclude residual granulocytes **(D)** that might have been inadvertently included in lymph-mono gate. CD66b$^{neg}$ cells were then gated for CD14$^{neg}$CD11c$^{pos}$ DCs **(E)**. CD66b$^{neg}$ cells were gated with CD14 and CD16 to exclude CD16$^{neg}$CD14$^{neg}$ cells **(F)**, and then further gated for HLA-DR$^{pos}$ to identify monocytes **(G)**. Dot plots were generated from representative HC sample using FlowJo. Plotted numbers depict frequency of cells from immediate parent gates. The gating strategy was used for Figures 3 and 4, and Supplemental Figures 3, 4, and 6.

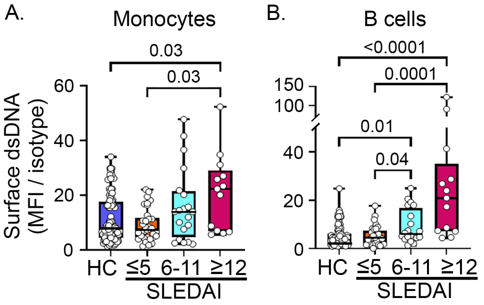


**Supplemental Figure 3. Accumulation of nuclear self-antigen on the surface of hematopoietic cells is not unique to nucleosome.** Unfractionated blood cells from HC (N= 101) or SLE patients (N=67) were stained with anti-dsDNA (33H11) or isotype control antibody, then analyzed by flow cytometry. N=13-101 patients per disease group, >10 experiments. Statistical analysis used Kruskal-Wallis. Adjusted $p$ values with significance are shown. Bar = median. Box= 25th-75th percentiles. Whiskers= minimum and maximum values.

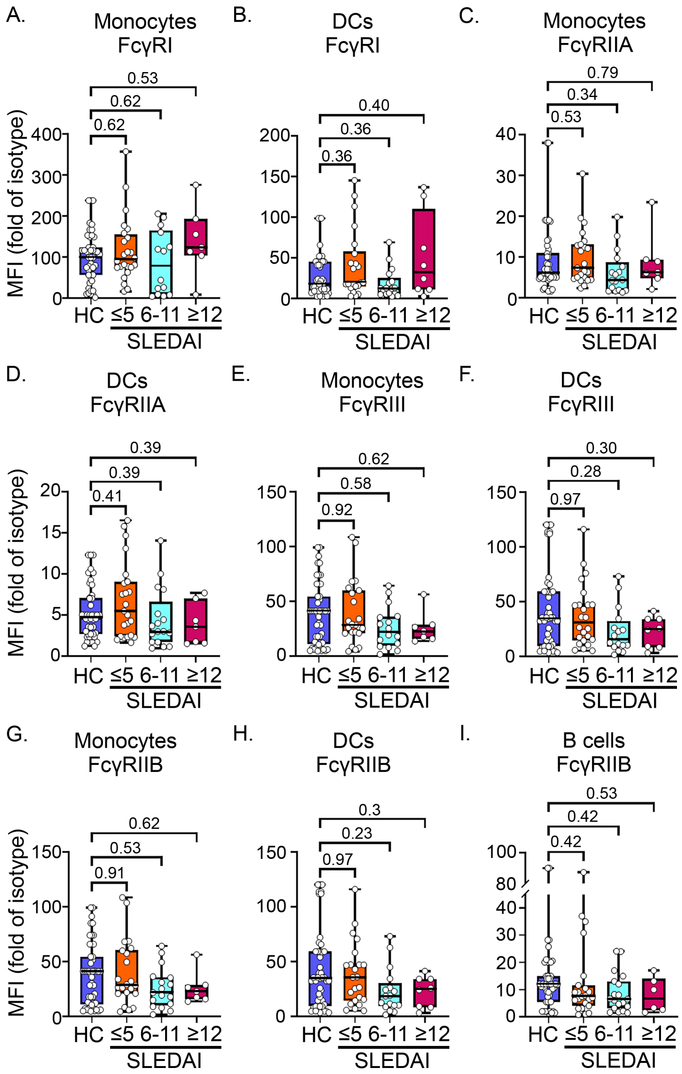


**Supplemental Figure 4. Levels of surface FcγRs are not different in patients, regardless of SLEDAI group. (A-I).** The levels of FcγRI, FcγRIIA, or FcγRIII on monocytes **(A, C, E)** and DCs **(B, D, F)** and FcγRIIb on monocytes **(G),** DCs **(H),** and B cells **(I)** were quantified by flow cytometry from unfractionated blood cells of HC or SLE patients. N = 6-48 patients per disease group, >6 experiments. Statistical analysis used Kruskal-Wallis. Adjusted *p* values are shown. Bar = median. Box= 25th-75th percentiles. Whiskers= minimum and maximum values.

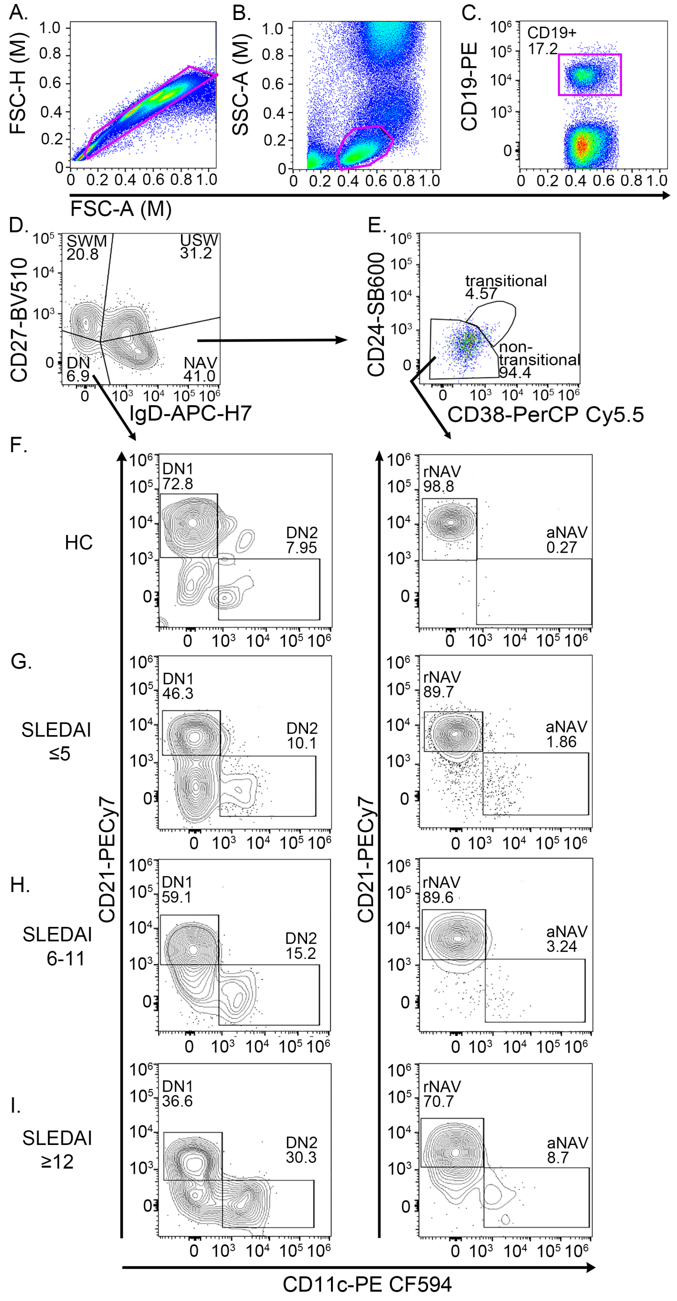


**Supplemental Figure 5. Flow cytometry gating for B cell subsets.** Unfractionated blood cells **(A)** were gated for lymphocytes **(B)**, then CD19+ B cells **(C)**. B cells were gated for switched memory (SWM), unswitched (USW), double-negative (DN), and naïve (NAV) cells using CD27 and IgD **(D)**. Naïve cells were sub-gated for transitional and non-transitional cells using CD24 and CD38 **(E)**. Non-transitional cells were further gated for resting naïve (rNAV) and activated naïve (aNAV) cells using CD21 and CD11c **(right panels in F-I)**. DN cells **(D)** were further gated for DN1 and DN2 cells using CD21 and CD11c **(left panels in F-I)**. Representative contour plots for DN1/DN2 and rNAV/aNAV cells from HC **(F)**, inactive **(G)**, moderately active **(H)**, and highly active **(I)** SLE are shown. Plots were generated with FlowJo. Gating strategy was used for Figure 4 **(D-G)**.

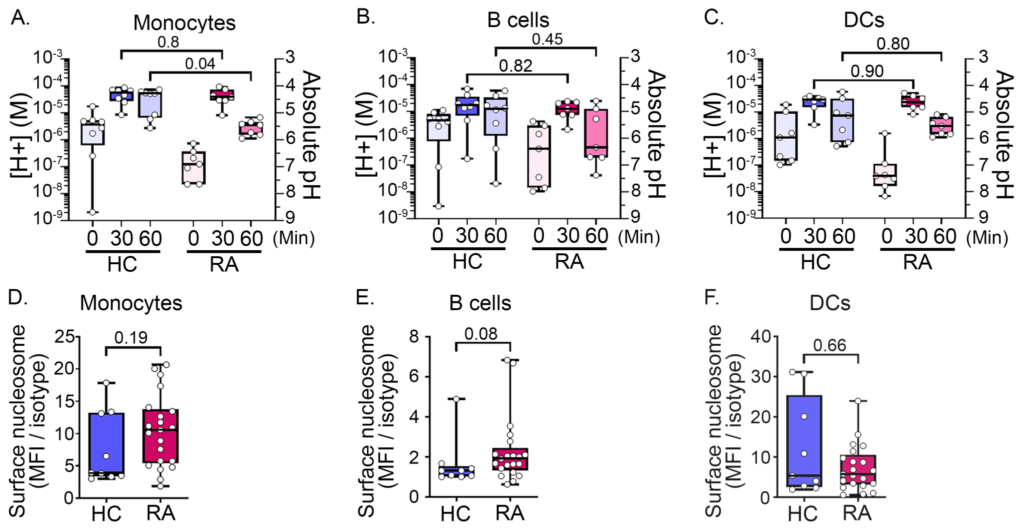


**Supplemental Figure 6. Late endosome/lysosome (LEL) dysfunction is not evident in rheumatoid arthritis (RA) patients.** LEL pH was measured in unfractionated blood cells from HC or active RA patients. Cells were stimulated with IgG-ICs (30 μl IgG-ICs/0.25x$10^6$ cells), then pH was measured by flow cytometry at designated time points **(A-C)**. To inhibit vATPase activity in unstimulated samples ($t_0$), cells were treated with Concanamycin A (20 ng/ml). Absolute pH was calculated using a standard curve, then converted to [H+] (pH = $-\log_{10}$ [H+]). Levels of surface nucleosome were quantified by flow cytometry **(D-F)**. N= 8 HC, 7 RA patients, 6 experiments **(A-C)**, N= 9 HC, 21 RA patients, >8 experiments **(D-F)**. Statistical analysis used Kruskal-Wallis **(A-C)** and Mann-Whitney **(D-F)**. Adjusted *p* values with significance are shown. Bar = median. Box= $25^{th}$-$75^{th}$ percentiles. Whiskers= minimum and maximum values.

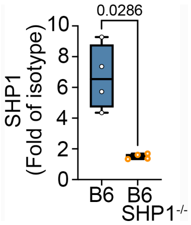


**Supplemental Figure 7. Tamoxifen treatment markedly reduced SHP1 in an inducible knockout model.** B6.SHP1$^{fl/fl}$ x B6. Rosa26-CreERT2 (B6.SHP1$^{-/-}$) mice were treated with 2mg of tamoxifen in 100 ul of corn oil per mouse (i.p.) for 4 consecutive days. Bone marrow cells were collected at day 17 post injection and cultured for 7 days to derive BMMφs. SHP1 levels in day 7 BMMφs from B6 and B6.SHP1$^{-/-}$ were measured by flow cytometry. N = 4, 2 experiments. Statistical analysis used Mann-Whitney test. Adjusted *p* values with significance are shown. Bar = median. Box= 25$^{th}$-75$^{th}$ percentiles. Whiskers= minimum and maximum values.

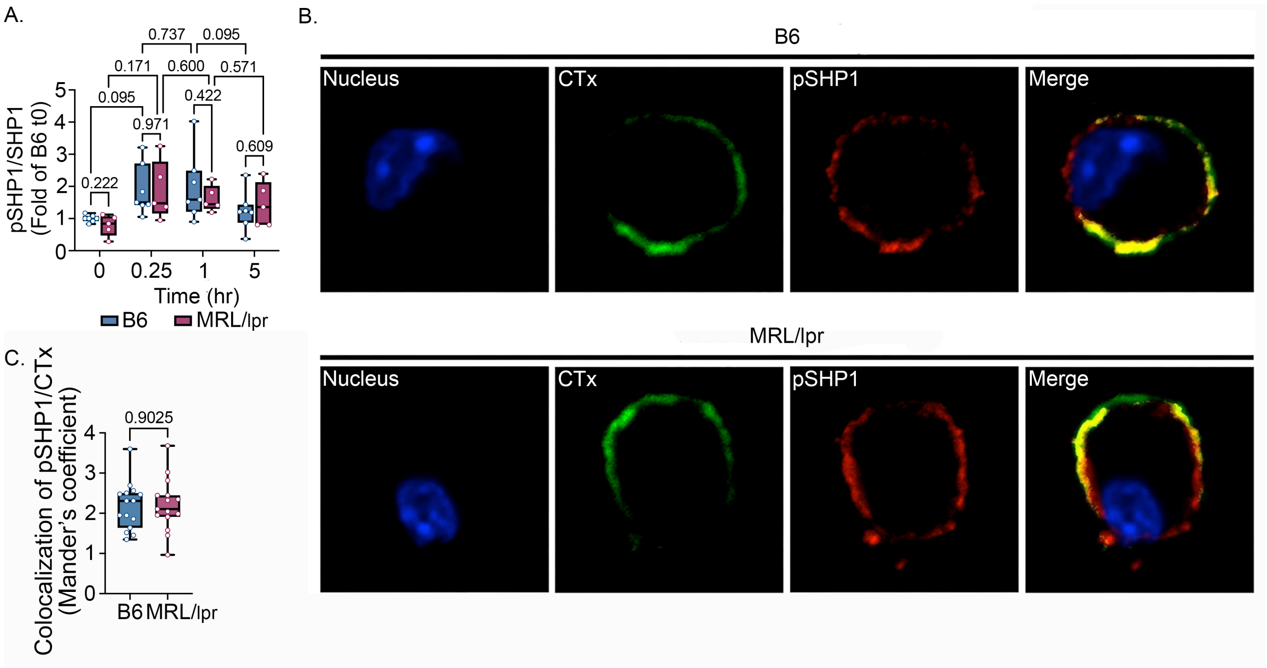


**Supplemental Figure 8. B6 and MRL/*lpr* mice showed comparable levels of pSHP1$^{Y564}$, and localization of pSHP1 to lipid raft.** BMMφs were treated with IgG-ICs (25μl IgG-ICs/0.25x10$^{6}$ cells) and at designated times, analyzed for intracellular pSHP1$^{Y564}$ levels by flow cytometry **(A)**. Fold of B6 $t_0$ was graphed. N= 5-7, 4 separate experiments. The localization of pSHP1$^{Y564}$ to plasma membrane lipid raft was assessed by confocal microscopy **(B)** and quantitated using Mander's coefficient **(C)**. BMMφs were treated with IgG-ICs (25μl IgG-ICs/0.25x10$^{6}$ cells) and stained for intracellular pSHP1$^{Y564}$ (red), lipid raft (cholera toxin, CTx, green), and cell nucleus (blue) **(B)**. Images were processed using Image J and representative images are shown. Yellow in merged images depicts colocalized pSHP1$^{Y564}$ and CTx. N=15 cells from 2 mice, 2 separate experiments. Statistical analysis used 2-way ANOVA with multiple comparisons **(A)** and Mann-Whitney test **(C)**. Adjusted *p* values with significance are shown. Bar = median. Box= 25$^{th}$-75$^{th}$ percentiles. Whiskers= minimum and maximum values.

#### Supplemental Tables

**Supplemental Table 1.** Median LEL pH and [H+] of each cell type in B6 and MRL/*lpr* mice of different ages.

| | B6 (pH[B]) | MRL/lpr (pH) | B6 ([H+] (M)[C]) | MRL/lpr ([H+] (M)) | Fold reduction over B6 ([H+] (M)) |
|---|---|---|---|---|---|
| CD11b+ myeloid cells[A] | | | | | |
| Age 9-10wks old | 4.5 | 4.7 | 2.84E-05 | 1.91E-05 | 1.5 |
| Age 12-13wks old | 4.4 | 5.1 | 4.26E-05 | 7.18E-06 | 5.9 |
| Age 15-16wks old | 4.3 | 5.4 | 4.48E-05 | 3.56E-06 | 12.6 |
| Age >18wks old | 4.7 | 5.7 | 1.97E-05 | 2.04E-06 | 9.7 |
| B cells | | | | | |
| Age 9-10wks old | 5.2 | 5.4 | 6.65E-06 | 4.46E-06 | 1.5 |
| Age 12-13wks old | 5.1 | 5.4 | 8.49E-06 | 3.83E-06 | 2.2 |
| Age 15-16wks old | 5.4 | 5.8 | 3.83E-06 | 1.66E-06 | 2.3 |
| Age >18wks old | 5.4 | 5.7 | 4.20E-06 | 2.03E-06 | 2.1 |
| DCs | | | | | |
| Age 9-10wks old | 5.1 | 5 | 7.67E-06 | 1.03E-05 | 0.7 |
| Age 12-13wks old | 5 | 5.5 | 1.12E-05 | 3.39E-06 | 3.3 |
| Age 15-16wks old | 5.4 | 5.5 | 4.32E-06 | 3.10E-06 | 1.4 |
| Age >18wks old | 5.2 | 5.6 | 6.75E-06 | 2.24E-06 | 3 |

[A]CD11b+ myeloid cells ($CD19^{neg}CD3^{neg}CD11b^{pos}CD11c^{neg/lo}$). [B] pH is calculated using a standard curve from cells stimulated with IgG-ICs for 30 min. [C] [H+] (M) is derived from absolute pH values using $[H^+] = 10^{-pH}$.

**Supplemental Table 2.** Demographic and clinical characteristics of participating SLE patients.

| | Overall |
|---|---|
| N | 81 |
| SLEDAI categories | |
| arthritis [N (%)] | 19 (23) |
| rash [N (%)] | 26 (32) |
| renal[A] [N (%)] | 24 (32) |
| vasculitis [N (%)] | 1 (1) |
| serositis[B] [N (%)] | 2 (3) |
| alopecia [N (%)] | 11 (13) |
| neuropsych[C] [N (%)] | 0 |
| ulcers [N (%)] | 5 (6) |
| hematologic[D] [N (%)] | 12 (15) |
| taking HCQ [N (%)] | 65 (80) |
| taking steroids [N (%)] | 33 (41) |
| taking DMARDs[E] [N (%)] | 46 (62) |
| Demographics | |
| age (mean ± SD[F]) | 40.4 ± 14 |
| Black [ N (%)] | 50 (65) |
| White [N (%)] | 27 (33) |
| Female [N (%)] | 72 (89) |
| Ethnicity | |
| Hispanic [N (%)] | 7 (9) |
| SLE History | |
| mean length of disease (yr) ± SD) | 11 ± 9.5 |
| historical renal disease[G] [N (%)] | 45 (56) |

[A]Renal: any SLEDAI renal parameter including active SLEDAI hematuria, pyuria, urinary casts, or proteinuria due to SLE. [B]Serositis: positive for pleurisy or pericarditis. [C]Neuropsych: positive for seizure, psychosis, organic brain syndrome, visual disturbance, cranial nerve disorder, lupus headache, or cerebrovascular accident. [D]Hematologic: positive for SLEDAI leukopenia or, thrombocytopenia. [E]DMARDS include Mycophenolic acid, Mycophenolate mofetil, Azathioprine, Methotrexate, and Tacrolimus. [F]SD = standard deviation. [G]Historic renal disease: ever had manifestation of nephritis based on ACR, SLICC, ACR/EULAR or renal SLEDAI.

**Supplemental Table 3.** Median LEL pH and [H+] in blood hematopoietic cells from HCs and SLE patients.

| | Mo | B cells | DCs |
|---|---|---|---|
| pH[A] | | | |
| HC | 4.8 | 5.3 | 4.8 |
| SLEDAI ≤ 5 | 4.8 | 5.4 | 4.7 |
| SLEDAI 6-11 | 5.4 | 5.8 | 5.5 |
| SLEDAI ≥12 | 5.6 | 5.9 | 5.7 |
| [H+] (M)[B] | | | |
| HC | 1.74E-05 | 5.62E-06 | 1.67E-05 |
| SLEDAI ≤ 5 | 1.80E-05 | 3.65E-06 | 2.19E-05 |
| SLEDAI 6-11 | 4.13E-06 | 1.76E-06 | 2.88E-06 |
| SLEDAI ≥12 | 2.66E-06 | 1.22E-06 | 2.01E-06 |
| [H+] (M) fold reduction/HC | | | |
| SLEDAI ≤ 5 | 1 | 1.5 | 0.8 |
| SLEDAI 6-11 | 4.2 | 3.2 | 5.8 |
| SLEDAI ≥12 | 6.5 | 4.6 | 8.3 |

[A]pH is calculated using a standard curve from cells stimulated with IgG-ICs for 30 min. [B][H+] (M) is derived from absolute pH values using $[H^+] = 10^{-pH}$.

**Supplemental Table 4.** Proportions of patients in each SLEDAI category showing non-acidic late endosomes/lysosomes (LELs), high surface nucleosomes, or high CIC levels for each cell type.

| | **arthritis** | **rash** | **renal**[D] | **alopecia** | **hematol**[E] | **HCQ** | **steroids** | **DMARDs**[F] |
|---|---|---|---|---|---|---|---|---|
| N | 19 | 26 | 24 | 11 | 12 | 33 | 46 | 12 |
| Low [H+] (Non-acidic) [N(%)][A] | | | | | | | | |
| Mo | 13 (68) | 19 (73) | 22 (92) | 7 (64) | 9 (75) | 23 (70) | 32 (70) | 9 (75) |
| B cells | 12 (63) | 19 (73) | 22 (92) | 8 (73) | 9 (75) | 24 (73) | 34 (74) | 9 (75) |
| DCs | 12 (63) | 17 (65) | 20 (83) | 8 (73) | 9 (75) | 23 (70) | 30 (65) | 9 (75) |
| High surface nucleosome [N(%)][B] | | | | | | | | |
| Mo | 1 (6) | 6 (29) | 5 (23) | 1 (14) | 3 (27) | 6 (22) | 7 (18) | 3 (27) |
| B cells | 2 (12) | 8 (38) | 11 (50) | 3 (43) | 2 (18) | 10 (37) | 14 (34) | 2 (18) |
| DCs | 1 (6) | 3 (14) | 4 (18) | 2 (29) | 3 (27) | 5 (19) | 9 (22) | 3 (27) |
| High CIC [N(%)][C] | | | | | | | | |
| CIC | 2 (50) | 9 (82) | 11 (85) | 3 (75) | 4 (50) | 12 (86) | 16 (76) | 4 (50) |

[A]Patients with “non-acidic” LELs have [H+] lower than the cut-off. For each cell type, the non-acidic cutoff was established at 1.8-fold above the mean [H+] of the HC. [B]Patients with “high surface nucleosome” have cell surface nucleosome levels above the cut-off. For each cell type, the cut-off was established at 1.8-fold above the mean surface nucleosome level of the HC. [C]Patients with “high CIC” show plasma CIC levels higher than the cut-off. The cut-off was established at 1.5-fold above the mean CIC level of the HC. [D]Renal – any SLEDAI renal parameter including active SLEDAI hematuria, pyuria, urinary casts, or proteinuria due to SLE. [E]Hematologic - positive for SLEDAI leukopenia or, thrombocytopenia. [F]DMARDS include Mycophenolic acid, Mycophenolate mofetil, Azathioprine, Methotrexate, and Tacrolimus. SLEDAI categories of vasculitis, serositis, and ulcers were not included because of few patients (1 vasculitis, 2 serositis, 5 ulcers).

**Supplemental Table 5.** Demographic and clinical characteristics of all participants with SLE and those with non-acidic late endosomes/lysosomes (LELs) for each cell type.

| | **Overall** | **Non-acidic[H] Monocytes** | **Non-acidic B cells** | **Non-acidic DCs** |
|---|---|---|---|---|
| N (%) | 81 | 53 (65) | 54 (67) | 46 (57) |
| SLEDAI categories | | | | |
| arthritis [N (%)] | 19 (23) | 13 (24) | 12 (22) | 12 (26) |
| rash [N (%)] | 26 (32) | 19 (36) | 19 (35) | 17 (37) |
| renal [N (%)][A] | 24 (32) | 22 (46) | 22 (44) | 20 (47) |
| vasculitis [N (%)] | 1 (1) | 1 (2) | 1 (2) | 1 (2) |
| serositis [N (%)][B] | 2 (3) | 2 (4) | 2 (4) | 1 (2) |
| alopecia [N (%)] | 11 (13) | 7 (13) | 8 (15) | 8 (17) |
| neuropsych [N (%)][C] | 0 | 0 | 0 | 0 |
| ulcers [N (%)] | 5 (6) | 2 (4) | 2 (4) | 3 (7) |
| hematologic [N (%)][D] | 12 (15) | 9 (17) | 9 (17) | 9 (20) |
| taking HCQ [N (%)] | 65 (80) | 42 (79) | 45 (83) | 36 (78) |
| taking steroids [N (%)] | 33 (41) | 23 (43) | 24 (44) | 23 (50) |
| taking DMARDs [N (%)][E] | 46 (62) | 32 (67) | 34 (72) | 30 (71) |
| Demographics | | | | |
| age (mean ± SD[F]) | 40.4 ± 14 | 38.2 ± 13.5 | 38.1 ± 13.5 | 44.9 ± 13 |
| Black [ N (%)] | 50 (65) | 34 (64) | 34 (68) | 28 (67) |
| White [N (%)] | 27 (33) | 15 (31) | 16 (32) | 14 (33) |
| Female [N (%)] | 72 (89) | 47 (89) | 47 (87) | 39 (85) |
| Ethnicity | | | | |
| Hispanic [N (%)] | 7 (9) | 6 (12) | 7 (14) | 5 (12) |
| SLE History | | | | |
| mean length of disease (yr) ± SD) | 11 ± 9.5 | 9 ± 7.3 | 10.9 ± 9 | 9.4 ± 7.9 |
| historical renal disease [N (%)][G] | 45 (56) | 33 (62) | 33 (61) | 29 (63) |

[A]Renal – any SLEDAI renal parameter including active SLEDAI hematuria, pyuria, urinary casts, or, proteinuria due to SLE. [B]Serositis - positive for pleurisy or pericarditis. [C]Neuropsych - positive for seizure, psychosis, organic brain syndrome, visual disturbance, cranial nerve disorder, lupus headache, or cerebrovascular accident. [D]Hematologic - positive for SLEDAI leukopenia or, thrombocytopenia. [E]DMARDS include Mycophenolic acid, Mycophenolate mofetil, Azathioprine, Methotrexate, and Tacrolimus. [F]SD = standard deviation. [G]Historic renal disease – ever had manifestation of nephritis based on ACR, SLICC, ACR/EULAR or renal SLEDAI. [H]Patients with “non-acidic” lysosomes have [H+] lower than the cut-off. For each cell type, the non-acidic cut-off was established at 1.8-fold above the mean [H+] of the HC.

**Supplemental Table 6.** Demographics of SLE patients analyzed for surface DNA.

| | Overall | SLEDAI ≤5 | SLEDAI 6-11 | SLEDAI ≥12 |
|---|---|---|---|---|
| N | 69 | 35 | 21 | 13 |
| Demographics | | | | |
| Age (mean ± SD[A]) | 42 ± 13 | 43 ± 13 | 44 ± 14 | 34 ± 8 |
| Black N (%) | 34 (49) | 21 (60) | 5 (24) | 8 (62) |
| White N (%) | 24 (35) | 11 (31) | 9 (43) | 4 (31) |
| Female N (%) | 61 (88) | 32 (92) | 20 (95) | 9 (69) |
| Ethnicity | | | | |
| Hispanic N (%) | 8 (11) | 2 (6) | 5 (24) | 1 (8) |

[A]SD = standard deviation. Missing data: race-1; ethnicity-4

**Supplemental Table 7**. Correlation between SLEDAI and molecular events associated with late endosome/lysosome (LEL) dysfunction.

| | **SLEDAI ≤5** | **SLEDAI 6-11** | **SLEDAI ≥12** |
|---|---|---|---|
| LEL [H+] Monocytes | | | |
| N | 44 | 24 | 13 |
| [H+] mean (SD[A]) | 2.7E-5 (2.8E-5) | 1.4E-5 (2.3E-5) | 8.6E-6 (1.3E-5) |
| Spearman's p-value | 0.84 | 0.69 | 0.56 |
| Correlation coefficient[B] | -0.03 | -0.09 | -0.18 |
| LEL [H+] B cells | | | |
| N | 44 | 24 | 13 |
| [H+] mean (SD) | 9.6E-6 (1.0E-6) | 7.1E-6 (1.0E-5) | 1.5E-6 (1.0E-6) |
| Spearman's p-value | 0.67 | 0.77 | 0.92 |
| Correlation coefficient | -0.07 | 0.06 | -0.03 |
| LEL [H+] DCs | | | |
| N | 44 | 24 | 13 |
| [H+] mean (SD) | 2.8E-5 (2.9E-5) | 9.7E-6 (1.5E-5) | 1.4E-5 (2.7E-5) |
| Spearman's p-value | 0.85 | 0.87 | 0.69 |
| Correlation coefficient | 0.03 | 0.03 | -0.12 |
| LEL hydrolase activity Monocytes | | | |
| N | 22 | 12 | 8 |
| MFI - mean (SD)[C] | 84.2 (67.7) | 53.3 (39.8) | 34.5(19.0) |
| Spearman's p-value | 0.75 | 0.4 | 0.91 |
| Correlation coefficient | 0.07 | 0.27 | -0.05 |
| LEL hydrolase activity B cells | | | |
| N | 20 | 11 | 8 |
| MFI - mean (SD) | 24.3 (29.7) | 12.3 (9.3) | 14.5 (11.8) |
| Spearman's p-value | 0.06 | 0.49 | 0.71 |
| Correlation coefficient | 0.49 | -0.23 | -0.16 |
| LEL hydrolase activity DCs | | | |
| N | 22 | 10 | 8 |
| MFI - mean (SD) | 43.3 (43.5) | 21.9 (14.2) | 21.2 (15.2) |
| Spearman's p-value | 0.54 | 0.7 | 0.98 |
| Correlation coefficient | 0.14 | -0.14 | -0.01 |
| Surface nucleosome Monocytes | | | |
| N | 37 | 24 | 10 |
| MFI - mean (SD) | 11.7 (10.2) | 12.8 (11.1) | 22.6 (15.8) |
| Spearman's p-value | 0.6 | 0.74 | 0.19 |
| Correlation coefficient | 0.09 | 0.07 | 0.45 |
| Surface nucleosome B cells | | | |
| N | 38 | 24 | 10 |
| MFI - mean (SD) | 7.8 (11.1) | 10.4 (10.4) | 32.5 (23.4) |
| Spearman's p-value | 0.55 | 0.26 | **0.004** |
| Correlation coefficient | 0.1 | 0.24 | **0.81** |
| Surface nucleosome DCs | | | |
| N | 38 | 24 | 10 |
| MFI - mean (SD | 8.3 (7.6) | 8.8 (7.2) | 12.4 (5.0) |
| Spearman's p-value | 0.29 | 0.91 | 0.36 |
| Correlation coefficient | -0.18 | -0.02 | 0.32 |
| Circulating immune complexes (CIC) | | | |
| N | 41 | 24 | 11 |
| mean (SD) | 7.1 (9.7) | 9.9 (9.4) | 22.4 (20.3) |
| Spearman's p-value | 0.68 | 0.42 | 0.68 |
| Correlation coefficient | 0.07 | 0.17 | 0.14 |

[A]SD = standard deviation. [B]Spearman's correlation coefficient and p-value. [C]MFI = mean fluorescence intensity

**Supplemental Table 8.** Demographics and clinical characteristics of SLE patients with high levels of CIC.

| | **Overall** | **high CIC**[H] |
|---|---|---|
| N (%) | 76 | 37 (49) |
| SLEDAI categories | | |
| arthritis [N (%)] | 18 (24) | 9 (24) |
| rash [N (%)] | 24 (32) | 13 (35) |
| renal [N (%)][A] | 22 (32) | 12 (36) |
| vasculitis [N (%)] | 1 (1) | 1 (3) |
| serositis [N (%)][B] | 1 (1) | 1 (3) |
| alopecia [N (%)] | 11 (15) | 6 (16) |
| neuropsych [N (%)][C] | 0 | 0 |
| ulcers [N (%)] | 5 (7) | 3 (8) |
| hematologic [N (%)][D] | 11 (15) | 6 (16) |
| plaquenil [N (%)] | 60 (79) | 30 (81) |
| steroids [N (%)] | 30 (40) | 16 (43) |
| DMARDs [N (%)][E] | 43 (61) | 20 (61) |
| Demographics | | |
| Age (mean ± SD[F]) | 41 ± 14 | 37 ± 13 |
| Black [N (%)] | 46 (64) | 22 (65) |
| White [N (%)] | 26 (36) | 12 (35) |
| Female [N (%)] | 68 (90) | 32 (87) |
| Ethnicity | | |
| Hispanic [N (%)] | 7 (10) | 7 (20) |
| SLE History | | |
| duration of disease in years (mean ± SD) | 10.5 ± 8.7 | 7.8 ± 4.9 |
| historical renal disease [N (%)][G] | 44 (58) | 23 (62) |

[A]Renal –any SLEDAI renal parameter including active SLEDAI hematuria, pyuria, urinary casts, or proteinuria due to SLE. [B]Serositis - positive for pleurisy or pericarditis. [C]Seuropsych - positive for seizure, psychosis, organic brain syndrome, visual disturbance, cranial nerve disorder, lupus headache, or cerebrovascular accident. [D]Hematologic - positive for SLEDAI leukopenia or, thrombocytopenia. [E]DMARDS include Mycophenolic acid, Mycophenolate mofetil, Azathioprine, Methotrexate, and Tacrolimus. [F]SD = standard deviation. [G]Historic renal disease – ever had manifestation of nephritis based on ACR, SLICC, ACR/EULAR criteria or renal SELENA-SLEDAI. [H]Patients with “high CIC” show plasma CIC levels higher than the cut-off. The cut-off was established at 1.5-fold above the mean CIC level of the HC.

**Supplemental Table 9.** Demographics of rheumatoid arthritis patients.

| | Overall |
|---|---|
| N | 23 |
| Demographics | |
| Age (mean ± SD[A]) | 53 ± 10 |
| Black N (%) | 7 (30) |
| White N (%) | 13 (57) |
| Other[B] | 3 (13) |
| Female N (%) | 18 (78) |
| Ethnicity | |
| Hispanic N (%) | 3 (13) |

[A]SD = standard deviation. [B]Other: Asian, Pacific Islander and not identified.

**Supplemental Table 10.** Proportion of patients in each renal disease activity group showing low [H+] in late endosomes/lysosomes (LELs) or high surface nucleosome.

| | Mo with non-acidic LEL[A] | B cells with non-acidic LEL | DCs with non-acidic LEL | Mo with high surface nucleosome[B] | B cells with high surface nucleosome | DCs with high surface nucleosome |
|---|---|---|---|---|---|---|
| Active Nephritis | | | | | | |
| N total | 25 | 25 | 25 | 23 | 23) | 23 |
| N positive (%) | 23 (92) | 22 (88) | 20 (80) | 5 (22) | 11(48) | 4 (17) |
| Remission Nephritis | | | | | | |
| N total | 20 | 20 | 20 | 15 | 15 | 15 |
| N positive (%) | 10 (50) | 11 (55) | 9 (45) | 3 (20) | 4 (27) | 2 (13) |
| Never Nephritis | | | | | | |
| N total | 36 | 36 | 36 | 33 | 33 | 33 |
| N positive (%) | 20 (54) | 21 (57) | 18 (49) | 3 (9) | 3 (9) | 6 (18) |

[A]Patients with "non-acidic" LELs have [H+] lower than the cut-off. For each cell type, the cut-off was established at 1.8-fold above the mean [H+] of the HC. [B]Patients with "high surface nucleosome" have cell surface nucleosome levels above the cut-off. For each cell type, the cut-off was established at 1.8-fold above the mean surface nucleosome level of the HC.
